# SPI-1 virulence gene expression modulates motility of *Salmonella* Typhimurium in a proton motive force- and adhesins-dependent manner

**DOI:** 10.1101/2023.02.01.526580

**Authors:** Doaa Osama Saleh, Julia A. Horstmann, María Giralt-Zúñiga, Willi Weber, Abilash Chakravarthy Durairaj, Enrico Klotzsch, Till Strowig, Marc Erhardt

## Abstract

Both the bacterial flagellum and the evolutionary related injectisome encoded on the *Salmonella* pathogenicity island 1 (SPI-1) play crucial roles during the infection cycle of *Salmonella* species. The interplay of both is highlighted by the complex cross-regulation that includes transcriptional control of the flagellar master regulatory operon *flhDC* by HilD, the master regulator of SPI-1 gene expression. Contrary to the HilD-dependent activation of flagellar gene expression, we report here that activation of HilD resulted in a dramatic loss of motility, which was dependent on the presence of SPI-1. Single cell analyses revealed that HilD-activation results in a SPI-1-dependent induction of the stringent response and a pronounced decrease of proton motive force (PMF), while flagellation was not affected. We further found that activation of HilD enhanced the adhesion of *Salmonella* to epithelial cells. A transcriptome analysis revealed a concomitant upregulation of several adhesin systems, which when overproduced, phenocopied the HilD-induced motility defect. We propose that a combination of SPI-1-dependent depletion of the PMF and upregulation of adhesins upon HilD-activation allows flagellated *Salmonella* to rapidly modulate their motility during infection, thereby enabling efficient adhesion to host cells and delivery of effector proteins.

## Introduction

*Salmonella enterica* is a rod-shaped, Gram-negative enteropathogen that causes human salmonellosis, including gastroenteritis and in rare cases enteric fever [1]. Important virulence factors include the flagellum and the injectisome, which is used to deliver effector proteins into host cells [2]. The flagellum is a rotating motility organelle that enables directed swimming towards nutrients and away from harmful substances [3]. Moreover, flagella are thought to contribute to *Salmonella* pathogenesis by enhancing the interaction with the host cell membranes and promoting actin polymerization [4,5]. Flagella are made of three main structures: (i) the basal body, embedded in the inner and outer membranes of *Salmonella*, (ii) a flexible hook, and (iii) a 10-15 µm long, helical filament [6]. The expression of flagella genes is regulated in a complex transcriptional hierarchy [7]. On top of the cascade resides the flagellar master operon *flhDC*. Although, six transcriptional start sites have been mapped within the promoter region of *flhDC*, only P1 and P5 were proved to be functional in a wild type background, where they drive its expression during early and late growth phases, respectively [8,9]. The transcription of *flhDC* is under the control of a σ^70^-dependent class I promoter and the protein products assemble in a heteromultimeric complex (FlhD_4_C_2_), which ultimately controls flagellar gene expression and flagellar assembly [9–11]. Functional FlhD_4_C_2_ directs the σ^70^/RNA polymerase holoenzyme to initiate transcription from class II promoters [12]. This results in the production of proteins needed for the construction of the basal body and hook, as well as regulatory proteins, such as the flagellar sigma factor FliA (σ^28^). FliA directs RNA polymerase to transcribe flagellar genes under the control of class III promoters. The products of those genes constitute the filament, the flagellar motor and chemosensory systems [13,14].

Injectisomes are specialized syringe-like needle complexes employed by *Salmonella* to inject effector proteins into eukaryotic host cells, thus manipulating them and facilitating the establishment of a successful infection [15,16]. The injectisome systems of *Salmonella* are encoded on the *Salmonella* Pathogenicity Islands 1 (SPI-1) and 2 (SPI-2) and play a role at different stages of the infection. The SPI-1-encoded injectisome is involved in the initial step of infection, where it translocates effector proteins into cells of the intestinal epithelium resulting in actin polymerization and remodelling [17,18]. This cytoskeletal remodelling results in ruffle formation on the host cell surface promoting the internalization of the bacterial cells. The SPI-2-encoded injectisome is crucial in the following steps, where it promotes trafficking of *Salmonella* across the epithelial cells as well as their replication and survival inside macrophages [19,20].

A battery of DNA-binding proteins, such as the AraC-like transcriptional regulators HilD, HilC and RtsA, regulate expression of SPI-1 genes. Each of those regulators can activate in a feed-forward loop its own transcription and the transcription of the other two regulators [21,22]. The three regulators can independently activate the expression of the OmpR/ToxR- type transcription factor HilA, which functions as the SPI-1 master regulator [23]. HilA directly activates the expression of SPI-1 injectisome structural components by binding to the SPI-1 encoded *prg*/*org* and *inv*/*spa* operons [24,25]. Additionally, it activates the expression of *invF*, an AraC-like transcriptional activator, as well as the SPI-1 chaperone *sicA*. InvF then interacts with SicA to induce the expression of effector proteins [26]. Both the flagellum and the injectisome are evolutionary related and, in particular, share a homologous protein export system, termed type-III secretion system [27,28]. To synthesize, assemble and secrete components of both systems, as well as to ensure flagellar rotation, significant amounts of energy are required [29]. Secretion of flagellar components is proton motive force (PMF)- dependent and coupled to ATP hydrolysis [30–33].

Moreover, *Salmonella* has the ability to rapidly adapt to multiple different conditions during the infection process. This requires a precise coordination of the production and operation of the injectisomes and the flagellum, which are both energy demanding processes. Several cross-regulatory connections between the different genetic systems that potentially contribute to this tight coordination process have been described in the literature. Kage et al. showed that the flagellar protein FliZ regulates HilD at the post-transcriptional level [34]. In contrast, expression of the flagellar master operon *flhDC* is influenced by various transcription factors, such as RflM/RcsB [35] or RtsB, which is transcribed in an operon together with the SPI-1 regulator RtsA [36]. RtsB binds within the *flhDC* promoter region to repress flagella biosynthesis. Additionally, FlhDC is post-transcriptionally regulated by the anti-FlhDC factor RflP (formerly known as YdiV) in response to nutrient levels and cell envelope stress [37,38]. HilD is also known to be involved in other regulatory networks in addition to its well-established role as an activator of SPI-1 gene expression [39]. In most of the so-far characterized cases, it exerts its regulatory function via relieving the repressive effect of the histone-like nucleoid-structuring protein (H-NS) on the promoters of the regulated genes [40–42]. For example, HilD activates the expression of SPI-2 genes via antagonizing the H-NS repression on the promoter region of the SPI-2 encoded *ssrAB* operon, favouring its OmpR-dependent expression [40,43]. SsrAB activates in turn the expression of SPI-2 genes. The expression of SPI-4 genes, which encode a PMF-dependent type-1 secretion system (T1SS) and its substrate protein SiiE, were also shown to be co-regulated with SPI-1 [44,45]. Further, SPI-5 encodes effectors that are translocated by the SPI-1-T3SS, are negatively regulated by H-NS and co-activated with SPI-1 genes under the same environmental conditions, which suggests that HilD is involved in their activation process [46,47]. In a previous study, we additionally described a transcriptional cross talk between SPI-1 and flagellar genes, where HilD directly binds within the promoter region of *flhDC* upstream of the P5 transcriptional start site and activates its expression [48].

In the present study, we report the surprising finding that activation of chromosomal HilD expression results in a pronounced motility defect in contrast to the increased flagellar gene expression upon *flhDC* activation. Transcriptome analysis revealed the HilD-dependent upregulation of genes encoding various adhesive structures such as chaperone-usher fimbriae or curli. Single cell analyses revealed that HilD-induced cells activated the stringent response and displayed a decrease in their membrane potential. Our results suggest that the observed motility defect upon HilD-activation is a multifaceted process that affects various cellular events. We propose that upregulation of adhesive structures and depletion of the PMF after activation of SPI-1 injectisome expression might allow *Salmonella* to rapidly adjust their motility behaviour during host cell infection independent of flagellar gene expression and assembly.

## Results

### Motility defect upon HilD-activation

We have previously shown that HilD directly activates flagellar gene expression by binding to the P5 transcriptional start site of the *flhDC* promoter [48]. However, the physiological relevance of the HilD-dependent activation of *flhDC* gene expression remained unclear. To characterize the HilD-dependent motility phenotype, we generated strains expressing a chromosomal copy of *hilD* under the control of an inducible promoter, including an ectopic locus (*araBAD*) and the native locus. HilD-dependent activation of gene expression was verified by monitoring the transcriptional activity of the promoter of the SPI-1 gene *sicA* fused to eGFP (P_*sicA*_-eGFP) and secretion of SPI-1-encoded effector proteins. As expected, transcription of *sicA* (Figure S1A) and secretion of effector proteins into the culture supernatant (Figure S1B) were both increased after HilD-induction. We next performed motility experiments to monitor the motility phenotype in response to HilD-activation. Surprisingly, swimming motility in soft-agar (0.3%) swim plates was drastically reduced (Figure 1A). Motility was restored to wildtype (WT) levels when a DNA binding-deficient mutant of HilD was expressed (*hilD*_∆HTH_). Overexpression of the transcriptional regulator HilA, which is downstream of HilD in the native SPI-1 regulatory cascade, also resulted in decreased motility. Since swimming motility in soft-agar swim plates is dependent on both bacterial growth and chemotaxis, we further assessed free-swimming motility of individual bacterial cells to exclude for those effects. We observed a pronounced decrease in the single-cell swimming velocities, which decreased to a mean speed of 1.5 µm/s after 45 minutes of HilD-induction (Figure 1B (right)), which coincided with increased transcription from the HilD-dependent promoter, P_*sicA*_. In the absence of HilD-induction (Figure 1B (left)), the mean swimming speed at that time point was 24 µm/s and the expression level of P_*sicA*_-eGFP was only 22.5 % of that in the HilD- induced cells. Moreover, after removing the inducer of *hilD* expression, swimming velocities were restored to WT levels (Figure S1C). Finally, in order to decouple production of flagella from possible HilD regulatory effects on the level of the *flhDC* promoter, we assessed motility in a strain in which the HilD binding site within the *flhDC* promoter region was randomized or deleted. Induction of HilD expression still resulted in non-motile bacteria, indicating that the observed motility phenotype was independent of a regulatory effect of HilD on *flhDC* transcription (Figure S1D).

**Figure 1:**
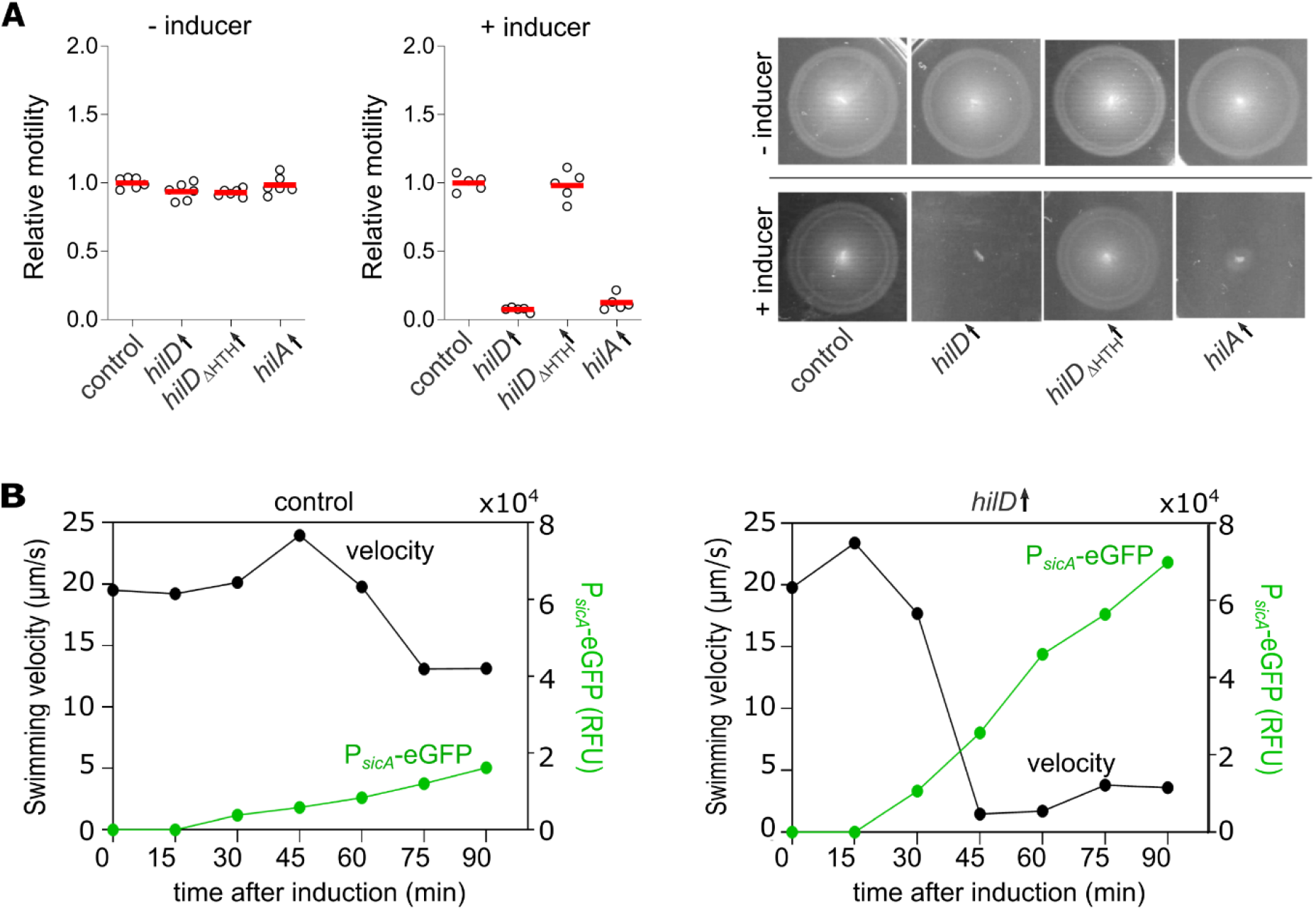
Activation of HilD abrogates swimming motility. **(A)** Swimming motility in soft-agar swim plates was monitored at 37 °C for 3.5 h in absence or presence of 0.2 % arabinose to induce HilD production. Diameters of swimming halos were measured and normalized to the control (left and middle panels). Representative swimming halos of the analysed mutants are shown (right). Biological replicates are shown as individual data points. Horizontal bars (red) represent the calculated mean of biological replicates. Statistical significances were determined using a two-tailed Student’s t-test (****, *P* < 0.0001; ns, *P* > 0.05). Strains analysed were EM808 (control), TH16339 (*hilD*↑), EM831 (*hilD*_ΔHTH_↑) and EM930 (*hilA*↑). **(B)** Induction of SPI-1 gene expression was monitored using a P_*sicA*_-eGFP transcriptional reporter fusion. Fluorescence intensities of P_*sicA*_-eGFP fusion and single-cell swimming velocities were measured after induction of HilD production by addition of AnTc (100 ng/ml). Fluorescence intensities of the cultures were measured in a microplate reader and normalised to the measured OD_600_ to give relative fluorescence units (RFU). The swimming velocities were analysed via time-lapse microscopy. Strains analysed were EM228 (control) and EM12302 (*hilD*↑). *hilD*↑: strain expressing *hilD* under an inducible promoter. *hilA*↑: strain expressing *hilA* under an inducible promoter. AnTc: anhydrotetracycline.

### Adhesion factors are involved in the HilD-mediated motility defect

We next performed transcriptome analysis to assess the impact of HilD-activation on *Salmonella* gene expression on a global scale. Principal component analysis (PCA) revealed that the transcriptome of the HilD-induced strain was distinct from the control strain (Figure S2A) and we were able to identify a large number of differentially expressed genes (> 2-fold change) under HilD-inducing conditions (Figure S2B). As expected, the expression levels of genes related to SPI-1 e.g. *prg, inv, org* and *spa* operons were increased under HilD-inducing conditions (Figure 2A). Additionally, genes of the SPI-2 operons (e.g. *sse* and *ssa*) and flagella genes (e.g., *flhD*) were upregulated. Interestingly, we also observed increased transcript levels of genes encoding for adhesive structures, such as SPI-4 genes, curli or other fimbrial operons (*pef* and *saf*). Moreover, the expressed levels of a previously described multiprotein immunoglobulin adhesion system ZirSTU, thought to be involved in an anti-virulence pathway, were elevated (Figure 2A). The upregulation of those genes was further validated using qRT-PCR (Figure 2B).

**Figure 2:**
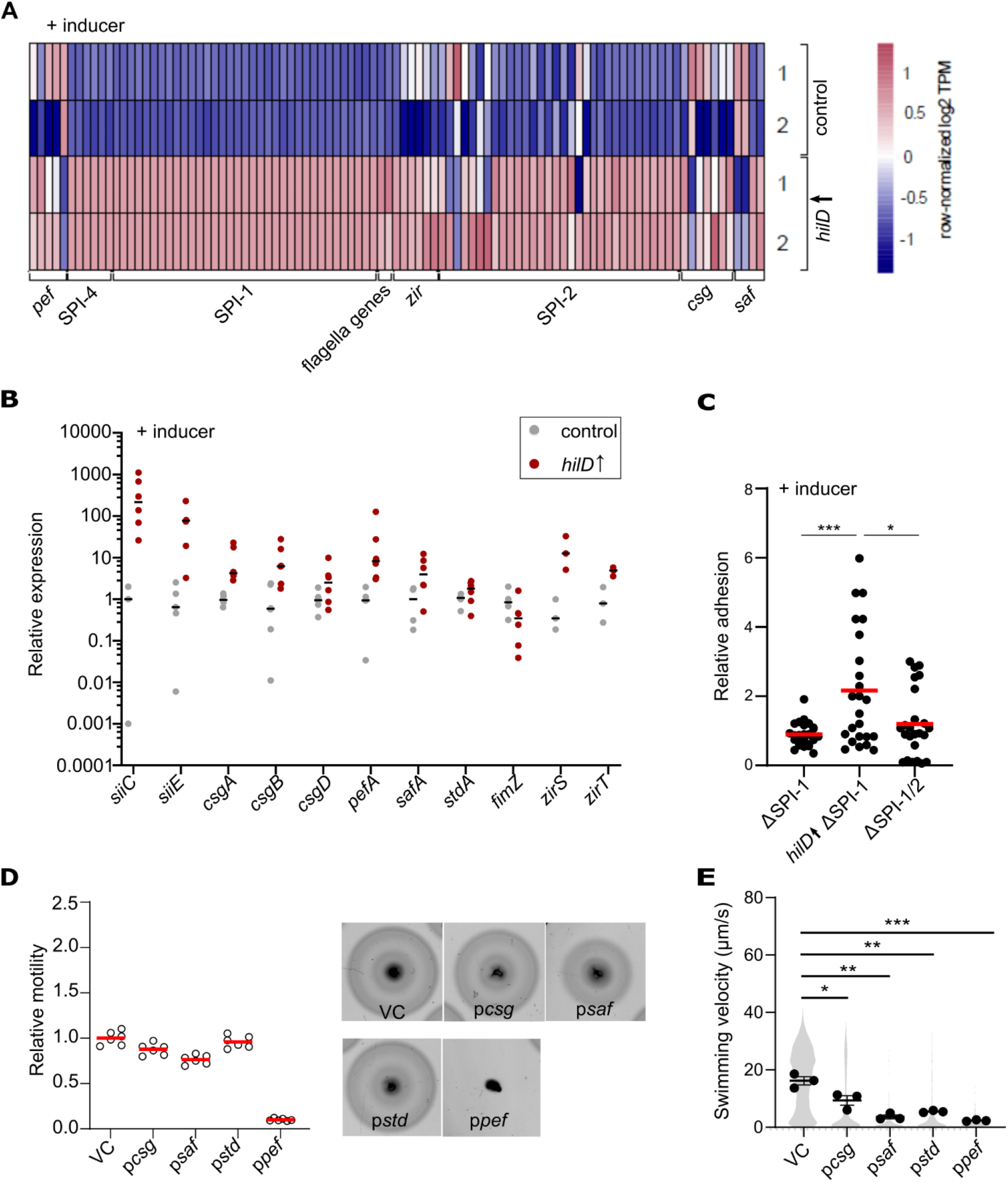
HilD activates expression of adhesive structures in *Salmonella* Typhimurium. (A) Heatmap of selected differentially expressed genes of the control (EM808) and *hilD*↑ (TH16339) strains. The TPM values of these selected genes were row normalised across the samples from two biological replicates (1, 2). (B) Validation of differential expression of *siiC, siiE, csgA, csgB, csgD, pefA, safA, stdA*, and *fimZ* in a HilD-induced strain compared to the control using qRT-PCR. (C) Relative adhesion of different *Salmonella* strains to MODE-K murine epithelial cells. MODE-K cells were incubated with different ΔSPI-1 strains at a MOI of 10 for 1 h at 37 °C. After extensive washing, cells were lysed and plated for CFU assessment. Counted CFU values were normalized to the ΔSPI-1 strain as a control. Biological replicates are shown as individual data points. Horizontal bars (red) represent the calculated mean of biological replicates. Statistical significances were determined using a two-tailed Student’s t-test (***, *P* < 0.001; *, *P* < 0.05). Strains analysed were EM830 (ΔSPI-1), EM93 (*hilD*↑ ΔSPI-1) and EM829 (ΔSPI-1/2). *hilD*↑: strain expressing *hilD* under an inducible promoter. (D) Swimming motility of strains overexpressing different adhesins in *trans* in soft-agar swim plates was monitored at 37 ° C for 3.5 h. Diameters of swimming halos were measured and normalized to the control strain (left). Representative swimming halos of the analysed mutants are shown (right). Biological replicates are shown as individual data points. Horizontal bars (red) represent the calculated mean of biological replicates. Strains analysed were EM12144 (VC, vector control), EM12145 (p*csg*), EM12146 (p*saf*), EM12147 (p*std*), EM12148 (p*pef*). (E) Single-cell swimming velocities of strains overexpressing different adhesins. Individual data points represent the average of the single-cell velocities of independent experiments. Violin plots represent all single-cell data values from all replicates. Horizontal bars (bold) represent the mean of the calculated average velocities of independent experiments. The error bars represent the standard error of mean and statistical significances were determined using a two-tailed Student’s t-test (***, *P* < 0.001; **, *P* < 0.01; *, *P* < 0.05). Strains analysed were the same as in panel (D). AnTc: anhydrotetracycline.

We therefore speculated that activation of HilD would induce expression of adhesive structures, which might in turn mediate the observed motility defect. We thus analysed adhesion of *Salmonella* to epithelial cells upon HilD induction in a strain deleted for SPI-1 to exclude for the increased adherence that might result from the enhanced docking conferred by the activated injectisomes. Additionally, epithelial cells were pre-treated with cytochalasin D to prevent actin polymerization and bacterial invasion. Strains expressing *hilD* under the native promoter and deleted for SPI-1 or both SPI-1 and -2 were used as control strains. As shown in Figure 2C, HilD-activation led to increased adhesion to the surface of MODE-K murine epithelial cells compared to the control strains. Accordingly, the contribution of fimbriae and curli to the HilD-dependent motility phenotype was assessed using strains overexpressing different fimbrial operons. On motility plates, only overproduction of Pef fimbriae resulted in a loss of motility, while Csg curli as well as Saf and Std fimbriae resulted in no or a very mild effect on motility (Figure 2D). In contrast, the overproduction of all tested fimbriae and curli constructs resulted in a sharp decrease of single-cell swimming velocities (Figure 2E). This suggested that the observed loss of motility in liquid medium might be due to increased adhesion to the glass surface of the flow chamber employed in the experimental setup, mediated by the overproduced adhesive structures.

### Deletion of SPI-1 restores motility upon HilD-induction

In addition to its identified role as an activator of *flhDC* expression, HilD is known to be involved in a broad regulatory network [39]. The role of HilD as a transcriptional activator of SPI-1 gene expression remains the best characterised [21,49,50]. Other targets include SPI- 2, SPI-4 and SPI-5 genes [37,39,40]. To investigate if the observed motility defect was dependent on any of those systems, we tested motility upon HilD-activation in genetic backgrounds deleted for each of them. Additionally, motility of a strain deleted for the genes encoding the twelve chaperone-usher fimbriae and the curli (Δ12 Δ*csg*) was tested. Interestingly, a ΔSPI-1 mutant displayed a WT motility behaviour upon HilD-induction on swimming motility plates (Figure 3A), as well as in liquid medium (Figure 3B). Deletions of other HilD-induced systems did not rescue the HilD-mediated motility decrease.

**Figure 3:**
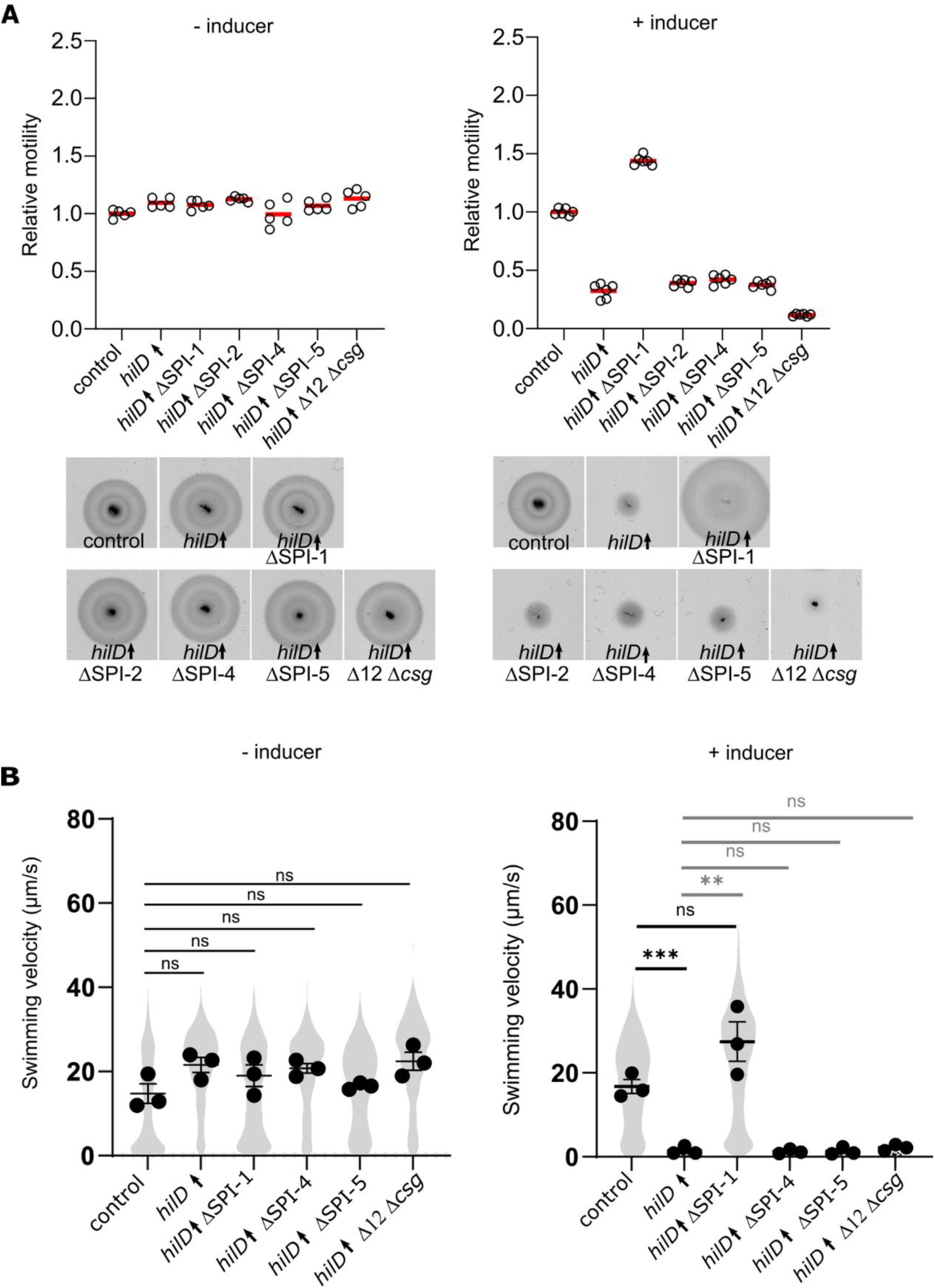
Deletion of SPI-1 restores motility upon HilD induction. (A) Swimming motility of the control, *hilD*↑ and various SPIs/adhesins mutant strains in soft-agar swim plates was monitored at 37 °C for 3.5 h in absence (upper panel) or presence (lower panel) of 100 ng/ml AnTc to induce HilD production. Diameters of swimming halos were measured and normalized to the control strain (left). Representative swimming halos of the analysed mutants in absence (upper panel) or presence (lower panel) of AnTc are shown (right). Biological replicates are shown as individual data points. Horizontal bars (red) represent the calculated mean of biological replicates. (B) Single-cell swimming velocities of the control, *hilD*↑ and various SPIs/adhesins mutant strains in absence or presence of AnTc to induce HilD production. Individual data points represent the average of the single- cell velocities of independent experiments. Violin plots represent all single-cell data values from all replicates. Horizontal bars (bold) represent the mean of the calculated average velocities of independent experiments. The error bars represent the standard error of mean and statistical significances were determined using a two-tailed Student’s t-test (***, *P* < 0.001; **, *P* < 0.01; ns, *P* > 0.05). Strains analysed were TH437 (control), TH17114 (*hilD*↑), EM12479 (*hilD*↑ ∆SPI-1), EM13020 (*hilD*↑ ΔSPI-2), EM12354 (*hilD*↑ ΔSPI-4), EM13021 (*hilD*↑ ΔSPI- 5) and EM12648 (*hilD*↑ Δ12 Δ*csgA*). *hilD*↑: strain expressing *hilD* under an inducible promoter. Δ12: strain deleted for 12 chaperone-usher fimbrial operons. AnTc: anhydrotetracycline.

### HilD induction activates the stringent response

We next investigated the growth kinetics of the bacteria upon HilD-activation using a microfluidic mother machine platform [52] (Figure 4A). Simultaneously, we determined the translational capacity of the cells by monitoring the fluorescence intensities resulting from the translation of a GFP expressed under an arabinose inducible promoter. Induction of HilD was associated with a growth defect and a decreased translation rate (Figure 4A, Figure S3A) as reported before [53]. Phase contrast microscopy revealed a reduction of the average cell length upon HilD-activation to ∼1.5 µm compared to ∼2.5 µm in WT cells, as well as a change in morphology from rod-shaped to coccoid (Figure S3B). Results from the mother machine platform confirmed that HilD-activation led to a ∼49% decrease in cell elongation rate (Figure 4B, Figure S3C (left)). The average maximal cell length during a division cycle decreased from 3.4 µm in the control cells to 2.2 µm in the HilD-induced cells. Consistently, the increase of cell length during a division cycle was ∼1.1 µm in HilD-induced cells compared to ∼1.5 µm in the control cells (Figure 4B, Figure S3C (middle and right)). The analysis further demonstrated an increase of the average generation time to ∼54 min upon HilD-induction compared to ∼28 min in the control (Figure 4B (left)). Although the mean GFP fluorescence intensity in the HilD-induced cells dropped to almost 47% of that of the control, the cells retained their translational capacity through the duration of the experiment (Figure 4A, Figure 4C (right)). Interestingly, deletion of SPI-1 was able to restore the growth rate defect, as well as cell length and morphology to the WT (Figure S3A, B). We therefore speculated that HilD-induction results in a cellular state that mimics a nutrient-limited environment. To investigate this possibility, we constructed a translational fusion of the fluorescent protein mCherry to *rpoS* in order to monitor activation of the stringent response. Expression of *rpoS* was enhanced in cells overproducing a hyperactive variant of the (p)ppGpp synthase RelA [54,55], validating the functionality of the stringent response reporter (Figure S4A). Additionally, *rpoS* levels were elevated when the cells were incubated in minimal medium devoid of carbon and nitrogen sources compared to their levels in the nutrient-rich LB medium (Figure S4B). We therefore investigated the levels of *rpoS* expression as a proxy for the stringent response in HilD-induced cells. As shown in Figure 4D, *rpoS* expression levels were elevated in HilD-induced cells compared to the control strain and decreased back to WT levels upon SPI-1 deletion, suggesting that HilD-activation induces the stringent response.

**Figure 4:**
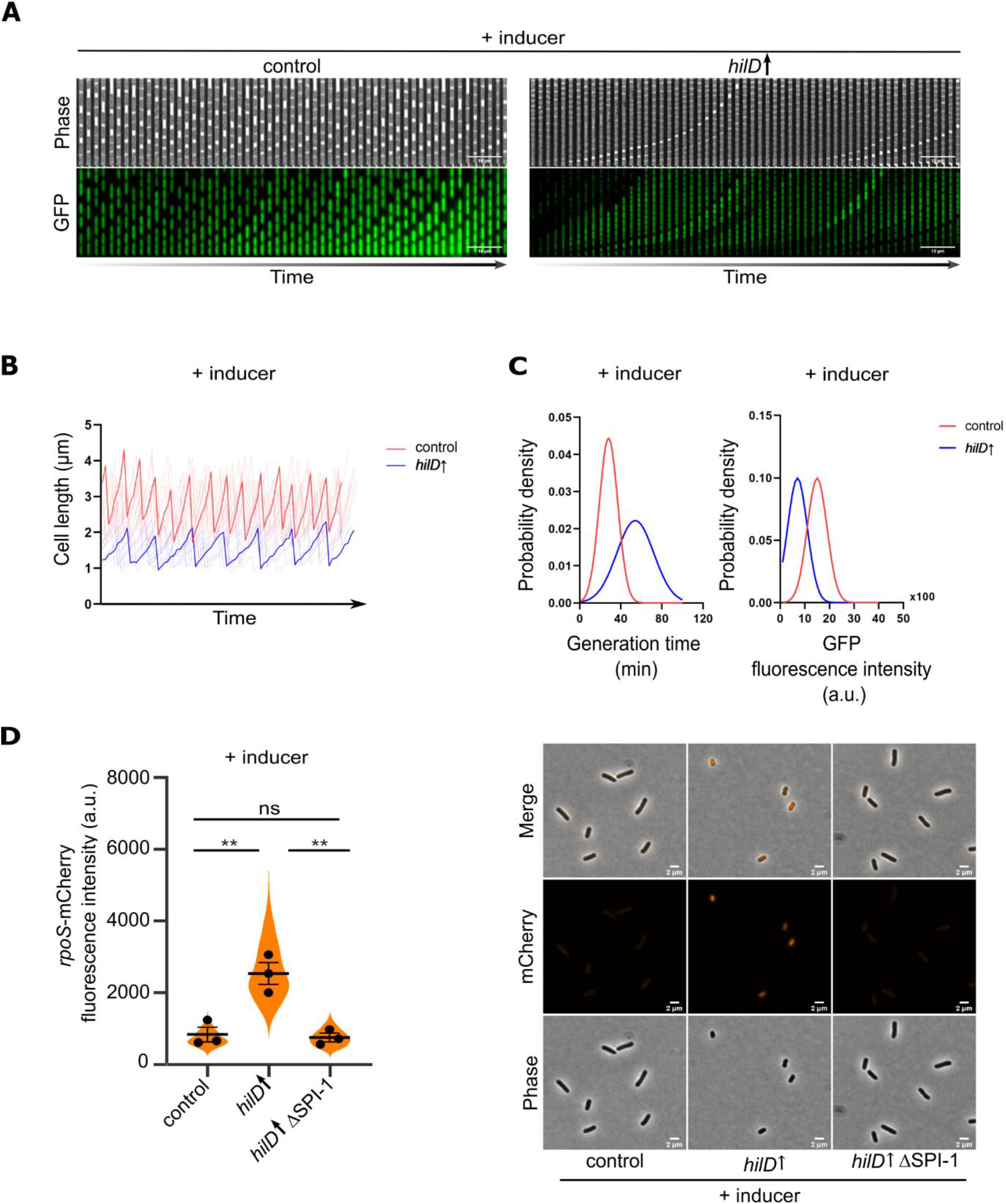
HilD induction activates the stringent response. (A) HilD-activation results in decreased growth rate and decreased translation rate. Strains were grown in the presence of AnTc as an inducer for HilD production. The expression of pBAD-GFP was achieved by addition of arabinose. The kymographs illustrate a single lineage of HilD-induced or control cells. Images from every second frame from the time-lapse were used to create the shown sequence. (B) Changes in cell length through time as determined using time-lapse microscopy for individual lineages. Traces of a sample of ten lineages are plotted in translucent colour and a sample trace is overlaid in opaque colour for each strain. (C) Probability density function of generation times (left) and GFP translation levels (right) in single cells as determined by time-lapse microscopy. Strains analysed in (A), (B) and (C) were EM12802 (control) and EM12803 (*hilD*↑). Violin plots represent single-cell data values from one individual experiment. *hilD*↑: strain expressing *hilD* under an inducible promoter. a.u.: arbitrary units. (D) HilD induction activates the expression of *rpoS*; a reporter for the stringent response. Fluorescence intensities of a *rpoS*- mCherry translational fusion were quantified using fluorescence microscopy (left). Individual data points represent the average of the single-cell fluorescence intensities from each independent experiments. Violin plots represent all single-cell data values from all replicates. Horizontal bars (bold) represent the mean of the calculated average velocities of independent experiments. The error bars represent the standard error of the mean and statistical significances were determined using a two-tailed Student’s t-test (**, *P* < 0.01; ns, *P* > 0.05). Representative microscopy images are shown (right). Scale bar is 2 µm. Strains analysed were EM13017 (control), EM13065 (*hilD*↑) and EM13276 (*hilD*↑ ΔSPI-1). *hilD*↑: strain expressing *hilD* under an inducible promoter. a.u.: arbitrary units.

### HilD induction is not associated with loss of flagellation

The anti-FlhDC factor, RflP (formerly known as YdiV) was shown before to tune the expression of flagella genes in response to nutrient levels [38]. Hence, we hypothesized that a nutrient-limited status after HilD-activation in addition to the general defect in translation result in enhanced *rflP* expression, which would act on FlhDC at the post-translational level catalysing its proteolytic degradation, thereby repressing flagella synthesis and motility. Subsequently, we investigated the levels of RflP expression using a translational fluorescent protein reporter fusion to *rflP*. As shown in Figure S5, we observed a small, yet statistically non-significant, increase in RflP expression upon HilD-activation, that was restored to WT levels in cells deleted for SPI-1. This result indicated that flagella synthesis might be impaired in response to the HilD-induced nutrient-limited cellular state. We therefore performed flagella immunostaining in order to evaluate the flagellation state upon HilD-activation. However, as shown in Figure 5, quantification of flagella numbers per cell demonstrated that also HilD- induced cells remained flagellated, albeit the average number of flagella per cell in HilD- induced cells was slightly decreased compared to the control strain.

**Figure 5:**
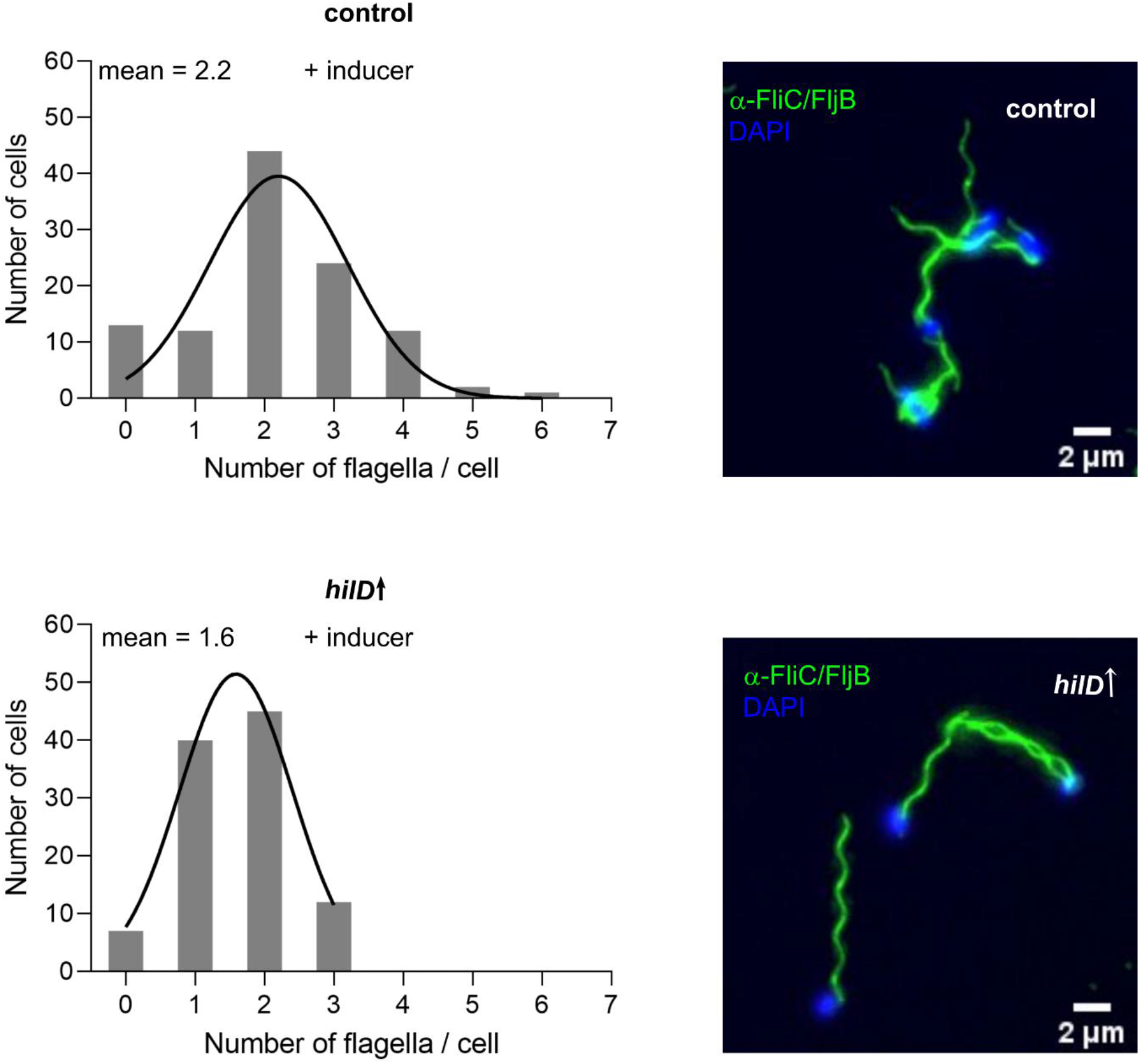
HilD induction does not affect flagellation. Counts of flagella per cell were determined using fluorescence microscopy. Histograms of counted flagella per cell are shown to the left. Average flagella numbers were calculated by Gaussian non-linear regression analysis (black line). Representative microscopy images of flagella immunostaining are shown to the right. Scale bar is 2 µm. Strains analysed were TH437 (control) and TH17114 (*hilD*↑). *hilD*↑: strain expressing *hilD* under an inducible promoter.

### Dissipation of the membrane potential contributes to the HilD-mediated motility defect

The proton motive force (PMF) serves as a main source of energy that drives secretion of protein substrates via the T3SS of both the flagellum and the injectisome [7,56]. Additionally, it energizes the rotation of the flagellar motor, thereby enabling swimming motility [57,58]. Therefore, we reasoned that increased secretion of injectisome substrates might dissipate the available PMF pool, thereby affecting flagella rotation and motility. Further, we recently reported that short-term starvation dissipates the PMF [59]. Similarly it was previously described by Verstraeten et al. that nutrient starvation induces a stringent response resulting in PMF dissipation in *Escherichia coli* [60]. Altogether, this suggests that induction of the stringent response observed upon HilD-activation might affect the membrane potential. We thus examined the membrane potential status in HilD-induced cells using the membrane potential-sensitive dye DiSC_3_(5) [59]. As a control, cells were treated with the ionophore CCCP prior to staining in order to dissipate the PMF. As expected, CCCP treated cells displayed diminished DiSC_3_(5) signal intensity (Figure S6). Interestingly, HilD-activation resulted in a sharp decrease of the membrane potential, which was largely restored upon SPI- 1 deletion (Figure 6). These results indicate that dissipation of PMF caused by increased assembly and/or activity of the injectisome and other PMF-draining systems might contribute to the observed motility defect.

**Figure 6:**
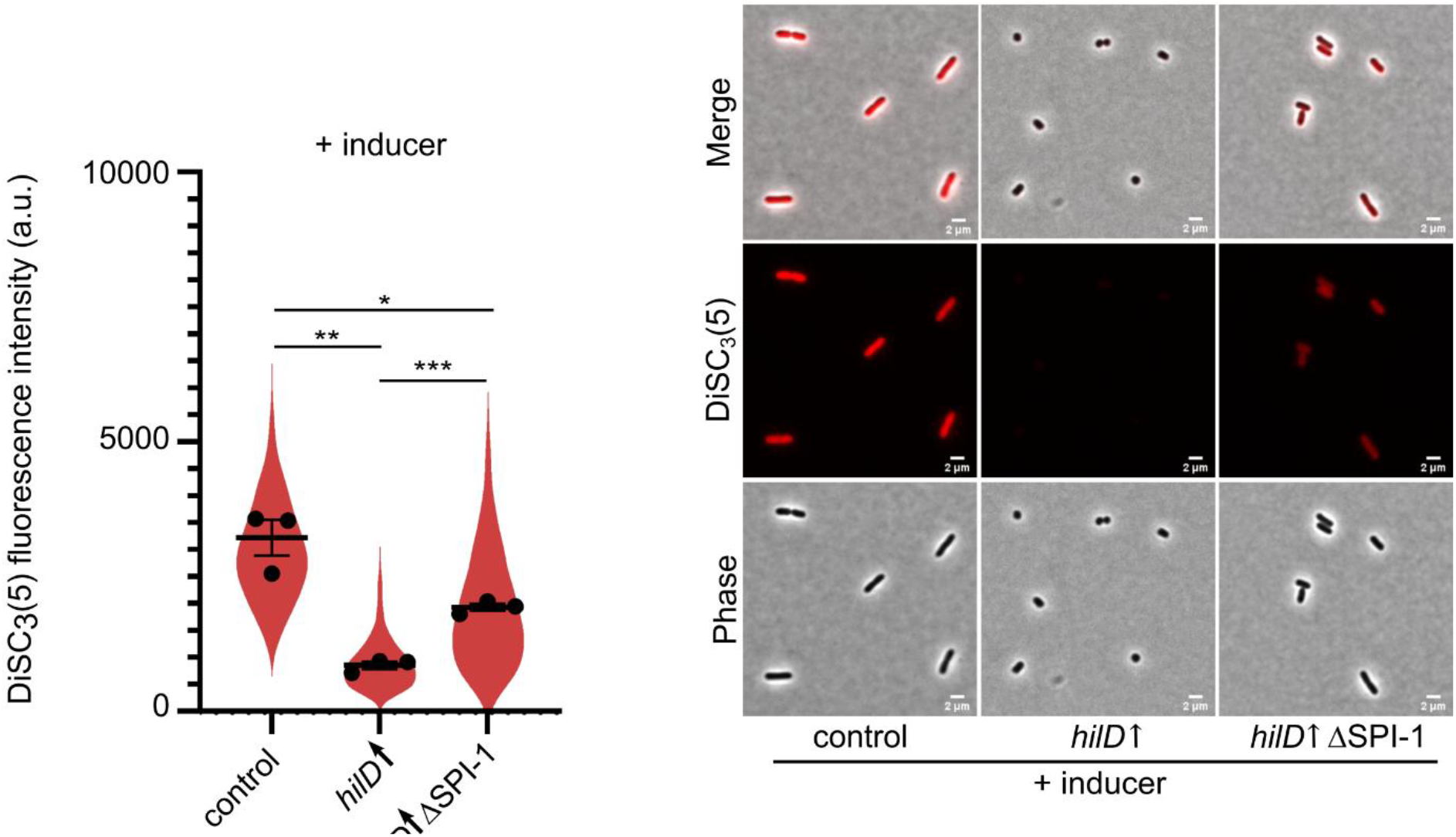
HilD induction results in membrane depolarization. Strains were grown under HilD-inducing conditions using AnTc followed by staining with the membrane potential sensitive dye DiSC_3_(5). DiSC_3_(5) fluorescence intensities of single cells were quantified (left). Individual data points represent the average of the single-cell values from each independent experiments. Violin plots represent all single-cell data values from all replicates. Horizontal bars (bold) represent the mean of the calculated average fluorescence intensities of independent experiments. The error bars represent the standard error of mean and statistical significances were determined using a two-tailed Student’s t-test. The error bars represent the standard error of the mean and statistical significances were determined using a two-tailed Student’s t-test (***, *P* < 0.001; **, *P* < 0.01; *, *P* < 0.05). Representative microscopy images are shown (right). Scale bar is 2 µm. Strains analysed were TH437 (control), TH17114 (*hilD*↑), EM12479 (*hilD*↑ ΔSPI-1), *hilD*↑: strain expressing *hilD* under an inducible promoter. AnTc: anhydrotetracycline. a.u.: arbitrary units.

## Discussion

Flagella-mediated motility is crucial for *Salmonella* to move efficiently through the host intestinal lumen and establish a successful infection. Production of flagella and the associated chemotaxis system requires expression of more than 60 genes [61] that are tightly regulated in response to the different environmental cues. Additionally, the flagellum is interconnected to the virulence-associated injectisome via a complex cross-regulatory network. A previously characterized crosstalk involves the transcriptional activation of the flagellar master regulatory operon *flhDC* by HilD, the master regulator of SPI-1. Here, we characterized the motility phenotype associated with the induction of HilD.

Unexpectedly, we observed a pronounced motility defect after HilD induction, while the cells surprisingly remained flagellated (Figure 1, Figure 5). We showed that motility was restored only in mutants lacking the SPI-1 locus, but not other SPIs. A transcriptome analysis upon HilD-induction revealed several upregulated genes that might contribute to the observed motility phenotype. As expected, genes related to SPI*-*1 and SPI-2 as well as SPI-4 were upregulated upon HilD induction. The methyl-accepting chemotaxis proteins, McpA and McpC, were also upregulated as described previously [62,63]. Moreover, various uncharacterized genes (STM05010, STM1329, STM1330, STM1600, STM1854, STM2585, STM4079s, STM4310, STM4312 and STM4313) were positively regulated by HilD, as described before by Smith et al. and Colgan et al. [62,64]. Interestingly, our transcriptome analysis revealed additional differentially regulated genes that were not described in previous studies [39,64], likely because of the prolonged time of HilD induction employed in the current study. Colgan et al. compared the transcriptomic profile of a *hilD* deletion strain to the WT, while Smith et al. used pulse expression of HilD from a plasmid in a *hilD* deletion mutant before analyzing the transcriptome compared to a deletion strain carrying an empty vector. While this approach enabled the identification of genes directly regulated by HilD, the longer HilD induction time used here enabled us to also identify genes that were indirectly affected by HilD presence. Accordingly, we could detect a number of upregulated genes that encode adhesive structures. (Figure 2). These adhesin-related genes include curli fimbriae genes encoded by the *csg* operons, the chaperone-usher fimbriae Pef and Saf as well as a multiprotein immunoglobulin adhesin system encoded by the *zirTSU* operon [45,55–65]. Consistent with the upregulations of adhesin operons, we observed an increased adhesion of *Salmonella* to intestinal epithelial cells upon HilD-activation (Figure 2C), which might explain the decreased swimming velocities of single cells (Figure 1B, Figure 2E). Further, overproduction of adhesin systems phenocopied the swimming defect of the HilD-induced strain. It is thus tempting to speculate that *Salmonella* induces both the SPI-1 encoded injectisome and several adhesin systems during the initial stage of infection in order to enhance binding of the bacteria to epithelial cells.

In line with previous studies, we additionally observed a growth defect upon HilD induction [53]. This growth defect was evidenced by a decrease in the cell elongation rate, the maximal observed cell length and an increased generation time. The growth retardation was associated on a cellular level with morphological changes from rod-shaped to coccoid. The deletion of SPI-1 was able to restore the growth rate and morphology of the cells. The morphological changes upon HilD-activation together with the observed decrease in the cell translational capacity suggested that the cells were undergoing a state of starvation [65–67]. This was further evidenced by the upregulation of *rpoS* expression (Figure 4), which has previously been used as a proxy for induction of the stringent response [68,69]. Although nutrient deprivation is known to activate the anti-FlhDC factor RflP [70], we found that the HilD-induced cells remained flagellated (Figure 5). Interestingly, a similar phenotype has previously been observed in *Rhizobium melitoli*, where starved cells lost motility while retaining normal flagellation [71]. Additionally, previous studies suggested that starvation conditions and induction of the stringent response result in dissipation of the membrane potential [59,60,72], which is critical for energizing flagellar rotation and motility. Recently, Sobota et al. observed a decrease in the PMF of cells expressing SPI-1 after exposure to external stress agents [73]. In agreement with these reports, we observed a decrease in membrane potential upon HilD-induction, which was restored by SPI-1 deletion (Figure 6).

In summary, we speculate that it might be important for *Salmonella* to quickly modulate its swimming motility upon reaching its target sites on the host cells in order to facilitate docking to the cells and to enable subsequent injection of effector proteins. Upon reaching the small intestine, *Salmonella* undergoes a phase of chemotaxis and flagellar motility inside the intestinal lumen in search for the intestinal epithelial cells [63]. Afterwards, it undergoes a process known as near-surface swimming, where its screens the epithelial cell surface for the permissive entry sites to start the invasion process [74,75]. In the absence of an active mechanism to eject already produced flagella as recently described for the polarly flagellated γ-proteobacteria [76], we propose a model where *Salmonella* employs a multi-faceted process upon HilD-mediated SPI-1 induction to upregulate adhesive structures and dissipate the membrane potential, therefore rapidly abrogating motility and priming the bacteria for efficient host cell invasion (Figure 7).

**Figure 7:**
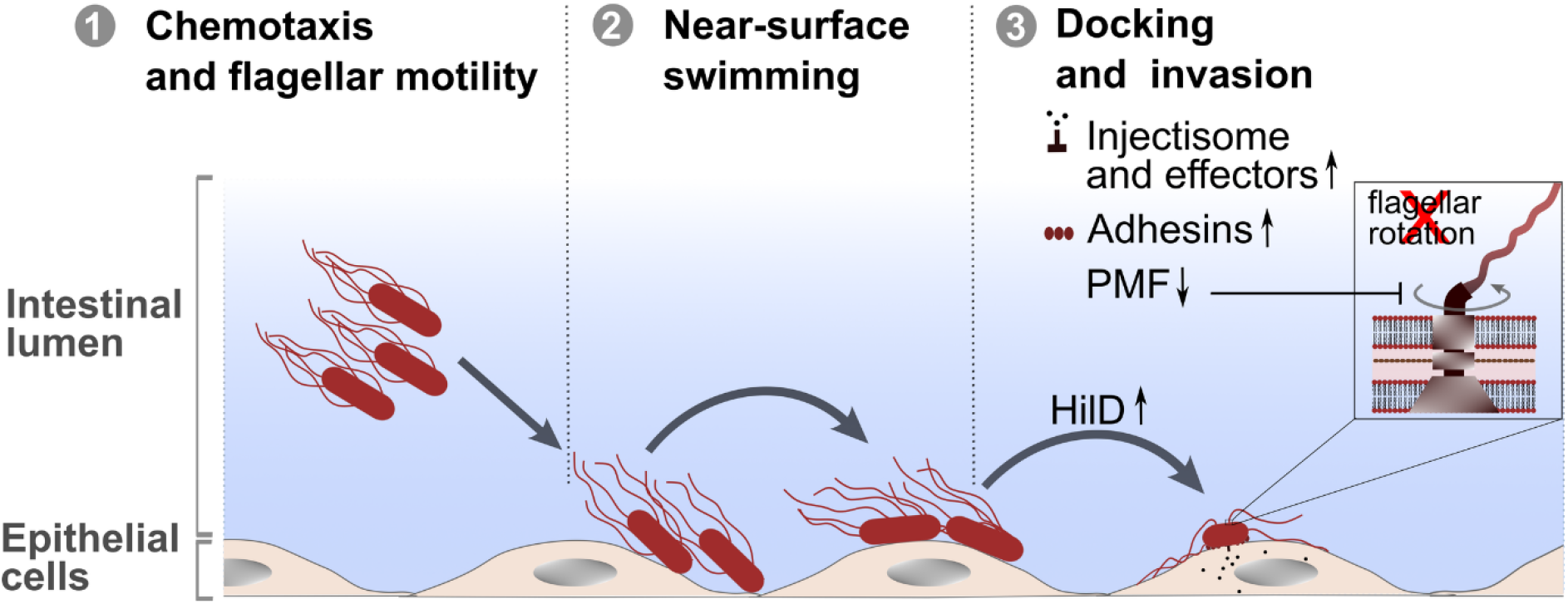
HilD-mediated SPI-1 induction abrogates motility. Inside the intestinal lumen, *Salmonella* goes through a phase of chemotaxis and flagellar motility in search for the host epithelial cells (1). Upon reaching the epithelial cells, *Salmonella* undergoes a phase of near-surface swimming in search for the permissive entry sites on the cells (2). After landing on the target site, HilD is upregulated to activate the SPI-1 encoded injectisome (3). Expression of SPI-1 and induction of HilD results in upregulation of adhesive structures and dissipates the membrane potential, therefore rapidly abrogating motility, enhancing docking to host cells and priming the bacteria for efficient host cell invasion.

## Materials and Methods

### Strains, media and bacterial growth

All *Salmonella enterica* serovar Typhimurium strains are listed in Supplementary Table S1. Bacteria were routinely grown in lysogeny broth (LB) at 37 °C, supplemented with 100 µg/ml ampicillin, 10 µg/ml chloramphenicol, 25 µg/ml kanamycin, 100 ng/ml anhydrotetracycline (AnTc) or 0.2 % arabinose when needed. Mutant strains were constructed using the general transducing *Salmonella* phage P22 HT105/1 int-201 [77] or using λ-RED recombination [78,79]. Bacterial growth was assessed via measurement of optical density at 600 nm (OD_600_) in a microplate reader (Tecan).

### Plasmid construction

To construct the plasmid expressing truncated RelA, the region coding for the N-terminal 455 amino acids of *relA* was PCR amplified from the genomic DNA of a wild type *Salmonella* strain (TH437) using cgc<u>catatg</u>GTCGCGGTAAGAAGTGCACA as a forward primer and ctag<u>tctaga</u>tcattattaCAACTGATAGGTGAATGGCA as a reverse primer. The primers were designed with overhangs (lowercase letters) containing restriction sites (underlined) for *Nde*I and *Xba*I enzymes, respectively. The digested product was cloned into the *Nde*I and *Xba*I sites on the pTrc99a-FF4 plasmid under the control of the IPTG-inducible *trc* promoter.

### Swimming motility

Swimming motility was assessed by inoculating 2 µl of overnight cultures of the desired strains into soft-agar swim plates containing 0.3 % agar followed by incubation at 37 °C for approximately 3.5 h.

### Single cell tracking

Bacteria were grown until mid-exponential growth phase in the presence of supplements where needed, then diluted into fresh LB to an OD600 ≤ 0.1. An aliquot was subsequently transferred to a flow chamber and a time-lapse video was recorded using a Nikon Eclipse Ti2 inverted microscope equipped with a CFI Plan Apochromat DM 20× Ph2/0.75 objective (Nikon) at 70 msec intervals. The obtained videos were segmented using the pixel classifier in Ilastik v1.3.3 [80] to recognize background and cells. Subsequently, all images in the dataset were processed to produce a “Simple Segmentation” mask output and exported to Fiji [81], where trajectories of the bacteria were tracked and the velocities of single cells were calculated using the simple LAP tracker of the Fiji plugin TrackMate [82].

### Fluorescence microscopy

For live cell microscopy, cultures were spotted on the surface of 1 % agarose pads (in PBS) cast on SuperFrost Plus™ slides (Erpedia). The spotted cultures were allowed to air-dry briefly, then covered with a microscopy cover slip (1.5H, Roth). For fixed cells imaging, in-house flow-chambers were constructed using slides and cover slips treated with 0.1 % poly-L-lysine (Sigma-Aldrich) [83]. The slide and cover slip were assembled in the presence of a double layered parafilm as a spacer. Samples were loaded into the chamber and allowed to adhere to the cover slip at room temperature (RT) followed by fixation with 4 % paraformaldehyde for 10 min. The fixed cells were then washed with PBS and mounted in Fluoroshield mounting medium containing DAPI (Sigma-Aldrich). Image acquisition was carried out using a Nikon Eclipse Ti2 inverted microscope equipped with a CFI Plan Apochromat DM 60× Lambda oil Ph3/1.40 objective (Nikon). The filter cube LED-CFP/YFP/mCherry-A (CFP / YFP / mCherry - Full Multiband Triple) (Semrock) was used for imaging *rpoS*-mCherry fusions and LED-DA/FI/TR/Cy5/Cy7-A (DAPI / FITC / TRITC / Cy5 / Cy7 - Full Multiband Penta) (Semrock) was used for P_*sicA*_-eGFP, *rflP*-mScarlet, DAPI and DiSC_3_(5).

### Microfluidic chip fabrication

Custom microfluidic master molds fabricated with Electron Beam Lithography were ordered from ConScience AB (Sweden). The mother machine chip used in this study features microchannels of 30 μm length, 1 μm width and 0.8 μm height. To prepare the PDMS chips, SYLGARD 184 silicone elastomer base (Dow, Europe) was mixed with the curing agent at a 7:1 ratio, degassed under vacuum for 30 minutes and poured onto the wafer and cured overnight at 80 °C. Inlets and outlets were punched in the chip using a disposable biopsy punch of 0.75 mm diameter. The PDMS chip was bonded to a high precision cover glass (24 × 60mm) using a 30 seconds oxygen plasma treatment in a plasma cleaner (Deiner) and incubated at 80 °C for 10 minutes.

### Microfluidic experimental setup and data analysis

Bacteria were grown in LB supplemented with ampicillin until late exponential phase. Cells were then loaded into the microchannels with a syringe tip connected to a syringe with Tygon tubing (inside diameter 0.51 mm and outside diameter 1.5 mm). LB supplemented with ampicillin, arabinose and AnTc was used as a growth medium and was delivered into the chip via Tygon tubbing connected to the inlets, using a syringe pump (World Precision Instruments). First the flow rate was set to 45 μl/min for 15 min to allow the main channels to clear, and then reduced to 5 μl/min for the duration of the experiment. The flow-throw was collected from the outlet with a second Tygon tubbing (inside diameter 0.51 mm and outside diameter 1.5 mm). Images were taken every 5 minutes for 20 hours. The automatic focalization was obtained via the Nikon PerfectFocus system. Bacmann software implemented in Fiji, was used for bacterial cell segmentation and tracking [81]. After initial pre-processing of the data (segmentation and rotation), centers of the microchannels were detected using the GFP channel, further allowing channel segmentation “*MicrochannelTracker* with *MicroChannelFluo2D* Segmenter”. Furthermore, the GFP channel was used to subsequently detect bacterial cells “*BacteriaClosedMicrochannelTrackerLocalCorrections* with *BacteriaFluo* module”. Time frames before the fluorescence signal reached a level sufficient for cell segmentation were excluded from further analysis. The resulting data were further analysed in Python 3.0.

### Flagella staining

Logarithmically grown cells were fixed on poly-L-lysine pre-coated coverslips as described previously followed by incubation with a solution of 10 % BSA to block non-specific antibody binding. The flagellar filament was immunostained by incubating with a 1:1 mixture of α-FliC and α-FljB primary antibodies (Difco) diluted 1:1,000 (in 2 % BSA solution). Samples were subsequently incubated with 10 % BSA to block the unspecific binding followed by incubation with a secondary α-rabbit antibody conjugated to Alexa Fluor 488 (Invitrogen) and diluted 1:1,000 (in PBS). Samples were mounted in Fluoroshield supplemented with the nucleic acid stain DAPI (Sigma-Aldrich).

### DiSC_3_(5) membrane potential assay

Logarithmically grown cultures were diluted to OD_600_ of 0.15 in LB medium. A volume of 500 µl was transferred to a 2 ml round bottom Eppendorf tubes, BSA to 0.5 mg/ml was added followed by the addition of the membrane potential sensitive dye 3,3’- Dipropylthiadicarbocyanine iodide (DiSC_3_(5)) [59]. The suspensions were allowed to incubate under shaking conditions in a thermomixer for 5 min. The lids of the Eppendorf tubes were left open to maintain sufficient aeration. Aliquots of 1 µl were then transferred to 1 % agarose pads (in PBS) and imaged immediately using Nikon Eclipse Ti2 inverted microscope where samples were excited at 647 nm for the detection of DiSC_3_(5) fluorescence. For validation of the DiSC_3_(5) dye, the diluted cultures were treated with the protonophore carbonyl cyanide m-chlorophenyl hydrazone (CCCP) at a final concentration of 1 mM for 15 min before adding the dye. Segmentation of bacterial cells and quantification of DiSC3(5) fluorescence intensities were performed using the Fiji plugin MicrobeJ [84].

### Microscopic evaluation of *rpoS*-mCherry reporter fusion

To validate the constructed *rpoS*-mCherry fusion in response to starvation conditions, a WT strain expressing the fusion was grown until mid-exponential phase under shaking conditions at 37°C in LB medium. Subsequently, the cells were centrifuged and resuspended in either M9 medium (Difco) supplemented with 1.0 mM CaCl_2_ and 0.1 mM MgSO_4_ but devoid of carbon or nitrogen sources or LB. The cells were then incubated for one additional hour. Aliquots were transferred to 1 % agarose pads for imaging. For validating the fusion in response to elevated levels of (p)ppGpp, a *Salmonella* strain carrying a plasmid expressing a truncated RelA comprised of the 455 N-terminal amino acids under an IPTG-inducible promoter was grown in LB supplemented with ampicillin under shaking conditions at 37 °C for approximately 2 h. Afterwards, IPTG was added to the culture to a final concentration of 0.2 mM and incubated for an additional hour. Subsequently, aliquots were transferred to 1 % agarose pads for imaging.

### Kinetic measurements of HilD-induced changes in single-cell swimming velocities and P_*sicA*_-eGFP expression

To monitor the effects of HilD induction on the swimming velocities and the expressed levels of the SPI-1 promoter fusion (P_*sicA*_-eGFP) in single bacterial cells, strains were grown at 37 °C to exponential phase in LB. Subsequently, 100 ng/ml anhydrotetracycline (AnTc) was added to induce HilD. Samples were taken immediately, and incubation of the cultures was resumed. Additional samples were then taken every 15 min. Single-cell swimming velocities were measured as described above and P_*sicA*_-eGFP fluorescence intensities were measured in the cultures using a microplate reader (Tecan). The measured fluorescence intensities were normalized to the OD_600_ of the cultures and reported as relative fluorescence units (RFU).

### Microscopic evaluation of HilD-activation

In order to validate the activity of HilD under control AnTc or arabinose inducible promoters, the levels of the HilD-dependent P_*sicA*_-eGPF transcriptional fusion were assessed as a reporter for SPI-1 genes activation. Strains expressing the P_*sicA*_-eGPF fusion and *hilD* under an inducible promoter were grown exponentially in absence or presence of 100 ng/ml AnTc or 0.2 % arabinose, respectively. Samples were transferred 1 % agarose pads and imaged as described above. Segmentation of bacterial cells and quantification of P_*sicA*_-eGPF fluorescence intensities were performed using the Fiji plugin, MicrobeJ [84].

### Adhesion assay

The murine epithelial cell line MODE-K was used for adhesion assays. 2.5 × 10^5^ cells/ml were seeded in 24-well plates and pre-treated with 1 µg/ml cytochalasin D for 30 min to inhibit actin polymerization. Cells were incubated with *Salmonella* strains at an MOI of 10 for 1 h. Afterwards the epithelial cells were washed extensively with PBS to remove non-adherent bacteria. Epithelial cells were subsequently lysed with 1 % Triton X-100 and cell lysates were serially diluted and plated on LB plates to determine the CFU/ml as a measure of bacteria that adhered to the epithelial cells. All values were normalized to the control strain.

### Secretion assay

*Salmonella* cultures were grown to mid-log phase in LB and then centrifuged at 4 °C to collect the culture supernatant. Secreted proteins were precipitated by addition of 10 % trichloroacetic acid (TCA, Sigma-Aldrich) followed by centrifugation for 1 h at 4 °C. The pellet was washed twice with acetone and air-dried. Samples were loaded and fractionated under denaturing conditions using SDS-PAGE. Proteins were subsequently stained using Coomassie Brilliant blue R250.

### Transcriptome analysis

Total bacterial RNA was isolated from logarithmically grown cultures by a hot phenol extraction protocol [85]. Residual DNA was digested using the TURBO DNase kit (Ambion) and RNA qualities were assessed using Agilent Technologies 2100 Bioanalyzer. Ribosomal RNA was depleted using Ribo-Zero rRNA Removal kit for bacteria (Epicenter). Library preparation was done using ScriptSeq kit (Epicenter). Sequencing was performed on HiSeq 2500 (Illumina) using TruSeq SBS Kit v3 - HS (Illumina) for 50 cycles. Image analysis and base calling were performed using the Illumina pipeline v 1.8.

### RNA isolation and quantitative real-time PCR

Total RNA was isolated from logarithmically grown cultures using RNeasy mini kit (Qiagen) as described previously [86]. Reverse transcription and quantitative real-time PCRs (qRT-PCR) were performed using the SensiFast SYBR No-ROX One Step kit (Bioline) in a Rotor-Gene Q lightcycler (Qiagen). Relative changes in mRNA levels were analysed according to Pfaffl [87] and normalized against the transcription levels of multiple reference genes according to the method described by Vandesompele et al. [88]. The reference genes *gyrB, gmk* and *rpoD* were used as previously described [48].

### Statistical analyses

Statistical analyses were performed with GraphPad Prism 9.0 (GraphPad Software, Inc., San Diego, CA) and values of *P* < 0.05 were considered statistically significant.

## Acknowledgements

We thank Michael Hensel (Universität Osnabrück) for kindly providing plasmids and Philipp F. Popp for his help with establishing single cell assays. We also thank Heidi Landmesser and Raúl Trepel for expert technical assistance. We are grateful for the Erhardt lab members for their valuable comments on the manuscript. This work was supported in part by a project that has received funding from the European Research Council (ERC) under the European Union’s Horizon 2020 research and innovation programme (grant agreement n° 864971) and from the Volkswagen (to M.E.) and by a GERLS scholarship (n° 91705821) co-funded by the Egyptian Ministry of Higher Education and Scientific Research (MHESR) and the German Academic Exchange Service (DAAD) (to D.O.S.). The funders had no role in study design, data collection and analysis, decision to publish, or preparation of the manuscript.

## Supporting information

**Figure S1:**
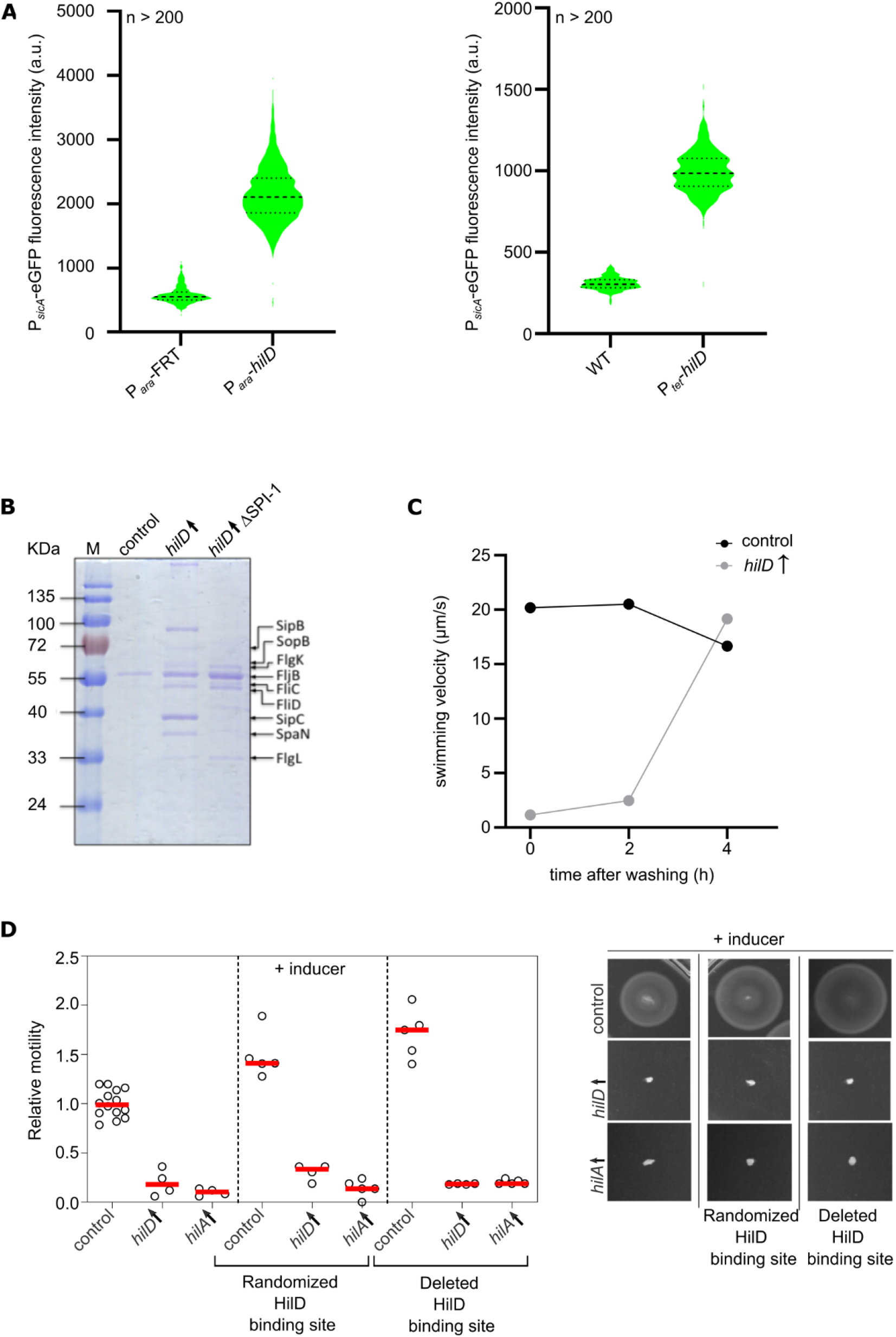
HilD induction increases SPI-1 effector protein transcription and secretion and affects motility independent of FlhDC. (A) Fluorescence intensities of a P_*sicA*_-eGFP fusion were measured in single cells as a reporter for *hilD* expression levels after inducing its expression from *araBAD* locus (P_*ara*_) (left) and from the native locus under a tetracycline inducible promoter (P_*tet*_) (right) using arabinose 0.2 % and AnTc 100 ng/ml, respectively. Strains analysed were EM899 (P_*ara*_-*hilD*), EM900 (P_*ara*_-FRT), EM228 (WT) and EM12302 (P_*tet*_- *hilD*). (B) SPI-1 effector protein secretion into the culture supernatant after HilD induction using arabinose 0.2 % was analysed via SDS-PAGE and Coomassie Blue staining. Strains analysed were EM808 (control), TH16339 (*hilD*↑) and EM93 (*hilD*↑ ΔSPI-1). (C) HilD-induced motility defect is reversed after removing AnTc used for inducing HilD production by washing with fresh medium. Swimming velocities of single cells were analysed via time-lapse microscopy in LB medium. Strains analysed were TH437 (control) and TH17114 (*hilD*↑). (D) Swimming motility in soft-agar swim plates was monitored at 37 °C for 3.5 h using strains mutated for HilD binding site in the *flhDC* promoter. Diameters of swimming halos were measured and normalized to the control (left). Representative swimming halos are shown (right). Biological replicates are shown as individual data points. Horizontal bars (red) represent the calculated mean of biological replicates. Strains analysed were EM808 (control), TH16339 (*hilD*↑), EM930 (*hilA*↑), EM3050 (control, randomized HilD binding site), EM3051 (*hilD*↑, randomized HilD binding site), EM3052 (*hilA*↑, randomized HilD binding site), EM3059 (control, deleted HilD binding site), EM3060 (*hilD*↑, deleted HilD binding site) and EM3061 (*hilA*↑, deleted HilD binding site).*hilD*↑: strain expressing *hilD* under an inducible promoter. *hilA*↑: strain expressing *hilA* under an inducible promoter. AnTc: anhydrotetracycline. a.u.: arbitrary units.

**Figure S2:**
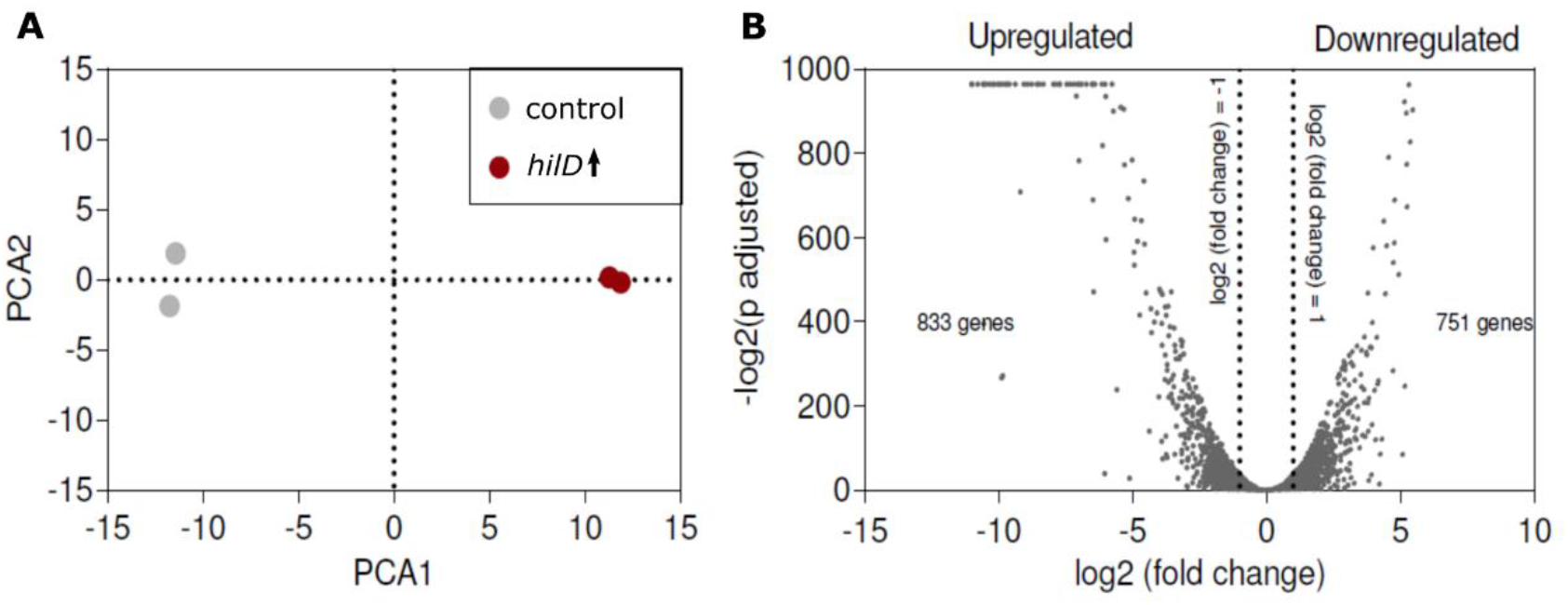
Transcriptome analysis upon HilD overexpression. (A) PCA plot for the control (EM808) and HilD-induced strains (TH16339) in the presence of arabinose as an inducer for HilD overproduction. Within-sample normalisation was done based on the Transcripts Per Million (TPM). The TPM values of genes less than one variance were removed. The log2 of the TPM values were used to make the PCA plot. (B) Volcano plot showing all differentially expressed genes. DESeq2 was used to calculate the log2(fold change) and the corresponding adjusted *p*-values for the genes, by comparing the expression profile of control samples against HilD-induced samples. *hilD*↑: strain expressing *hilD* under an inducible promoter.

**Figure S3:**
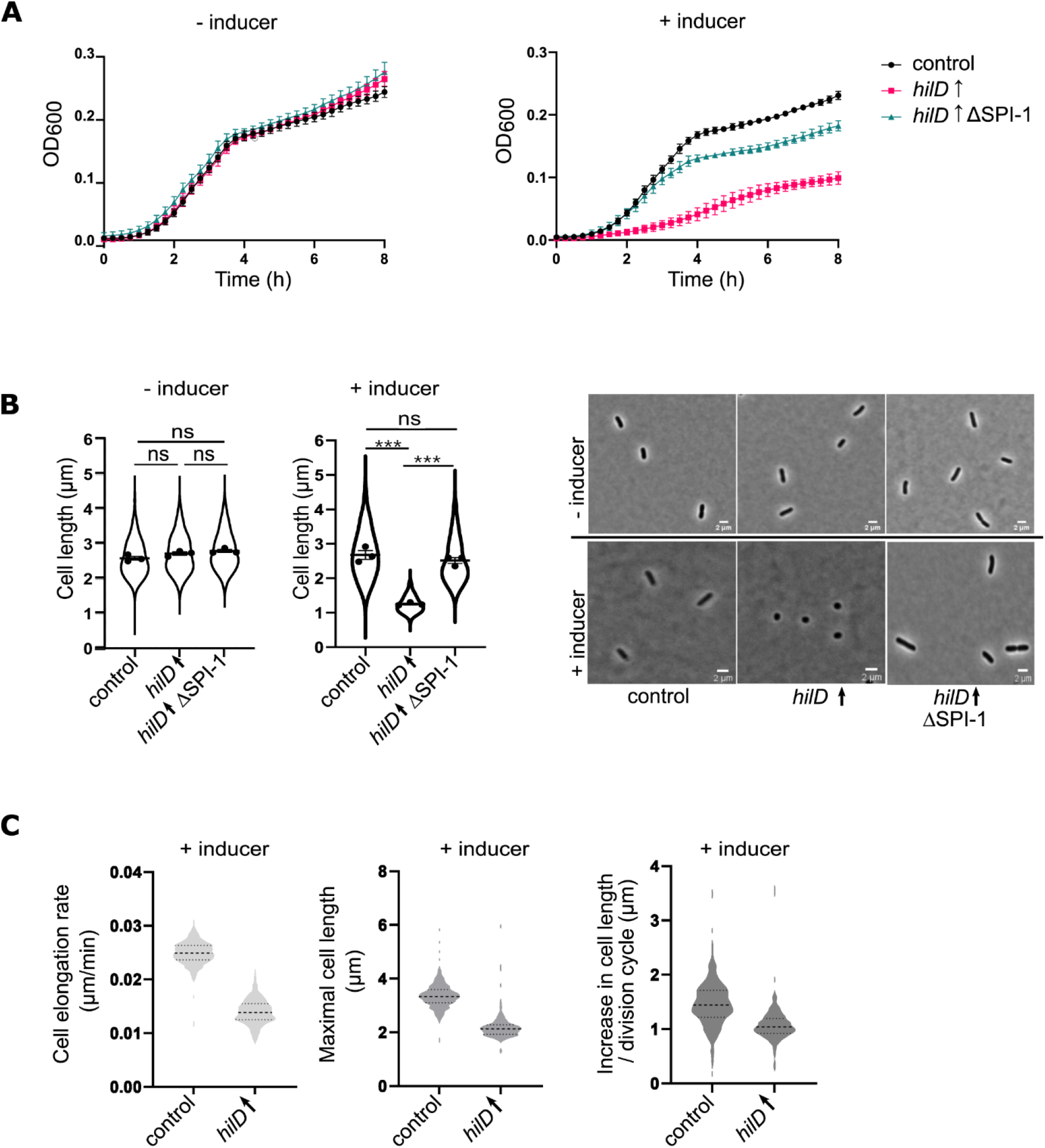
HilD-activation results in a growth defect. (A) Growth rates determined as a function of optical density at 600 nm (OD_600_) in absence and presence of AnTc for inducing HilD production. Individual points represent means of biological replicates at each time point. Error bars represent the standard deviation. *hilD*↑: strain expressing *hilD* under an inducible promoter. AnTc: anhydrotetracycline. Strains analysed were TH437 (control), TH17114 (*hilD*↑), EM12479 (*hilD*↑ ΔSPI-1). (B) HilD induction results in decreased cell length (left) and coccoid morphology (right). Cell length (µm) of individual bacteria was determined by phase-contrast microscopy. Individual data points represent the average of the single-cell lengths of independent experiments. Violin plots represent all single cell data values from all replicates. Horizontal bars (bold) represent the mean of the calculated average lengths of independent experiments. The error bars represent the standard error of mean and statistical significances were determined using a two-tailed Student’s t-test (***, *P* < 0.001; ns, *P* > 0.05). Representative images are shown in the right panel. the same strains as in (A) were analysed. (C) Time-lapse microscopy analysis reveals a decreased cell elongation rate (left), a decrease in the maximal cell length (middle) as well as decreased length elongation during a division cycle (right) upon HilD induction. Strains analysed were EM12802 (control) and EM12803 (*hilD*↑).

**Figure S4:**
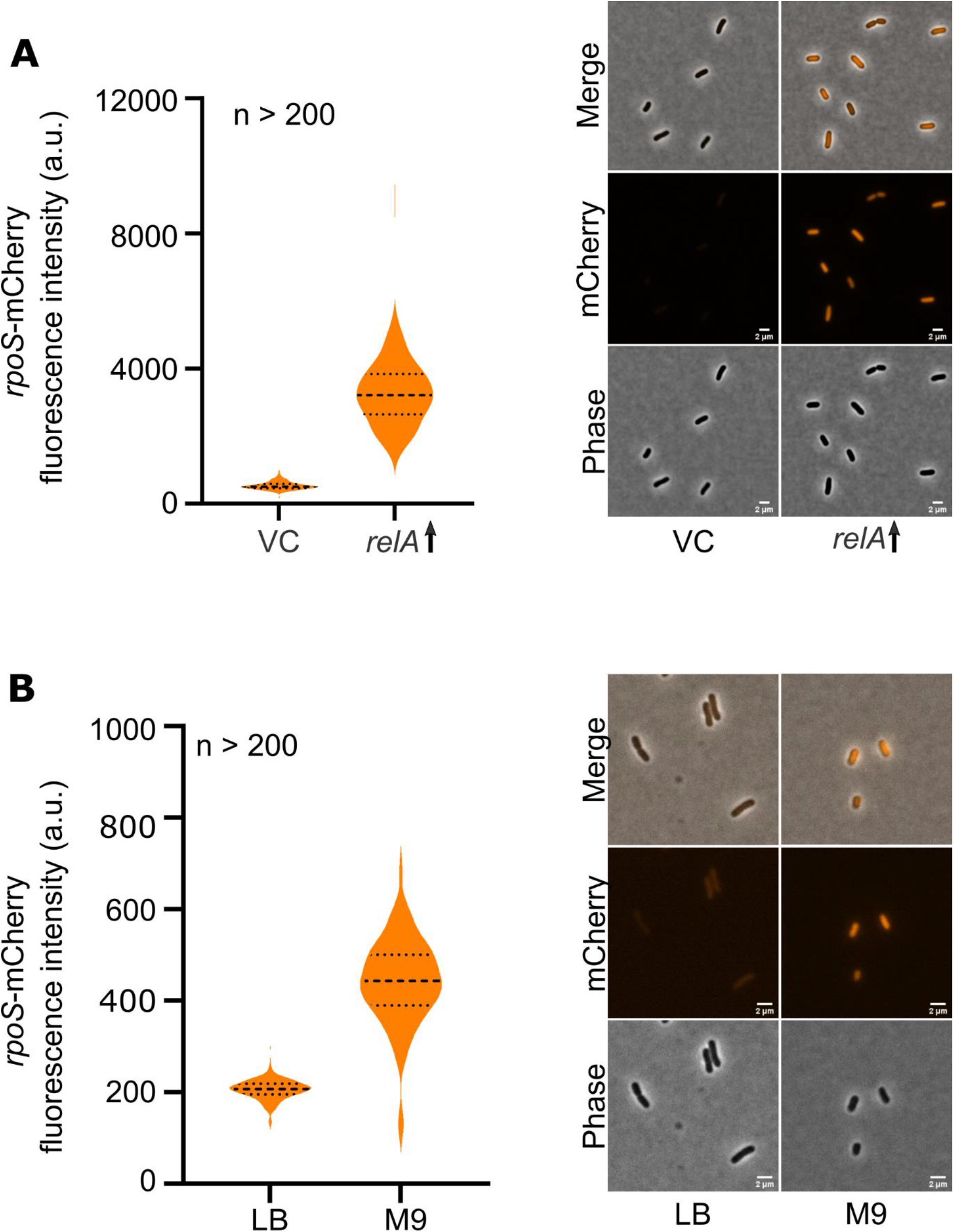
*rpoS*-mCherry functions as a reporter for the stringent response. (A) Fluorescence intensities of *rpoS*-mCherry C-terminal translation fusions were measured in a strain overexpressing a constitutively active RelA mutant comprised of the first N-terminal 455 amino acids (*relA*↑). Fluorescence intensities in single cells were quantified using fluorescence microscopy (left). Representative images are shown (right). Strains analysed were EM13226 (VC) and EM13227 (*relA*↑). (B) Assessment of *rpoS*-mCherry response to nutrient limitation in M9 minimal medium. Fluorescence intensities in single cells were quantified using fluorescence microscopy (left). Representative images are shown (right). Scale bar is 2 µm. Strain analysed was EM13017. a.u.: arbitrary units.

**Figure S5:**
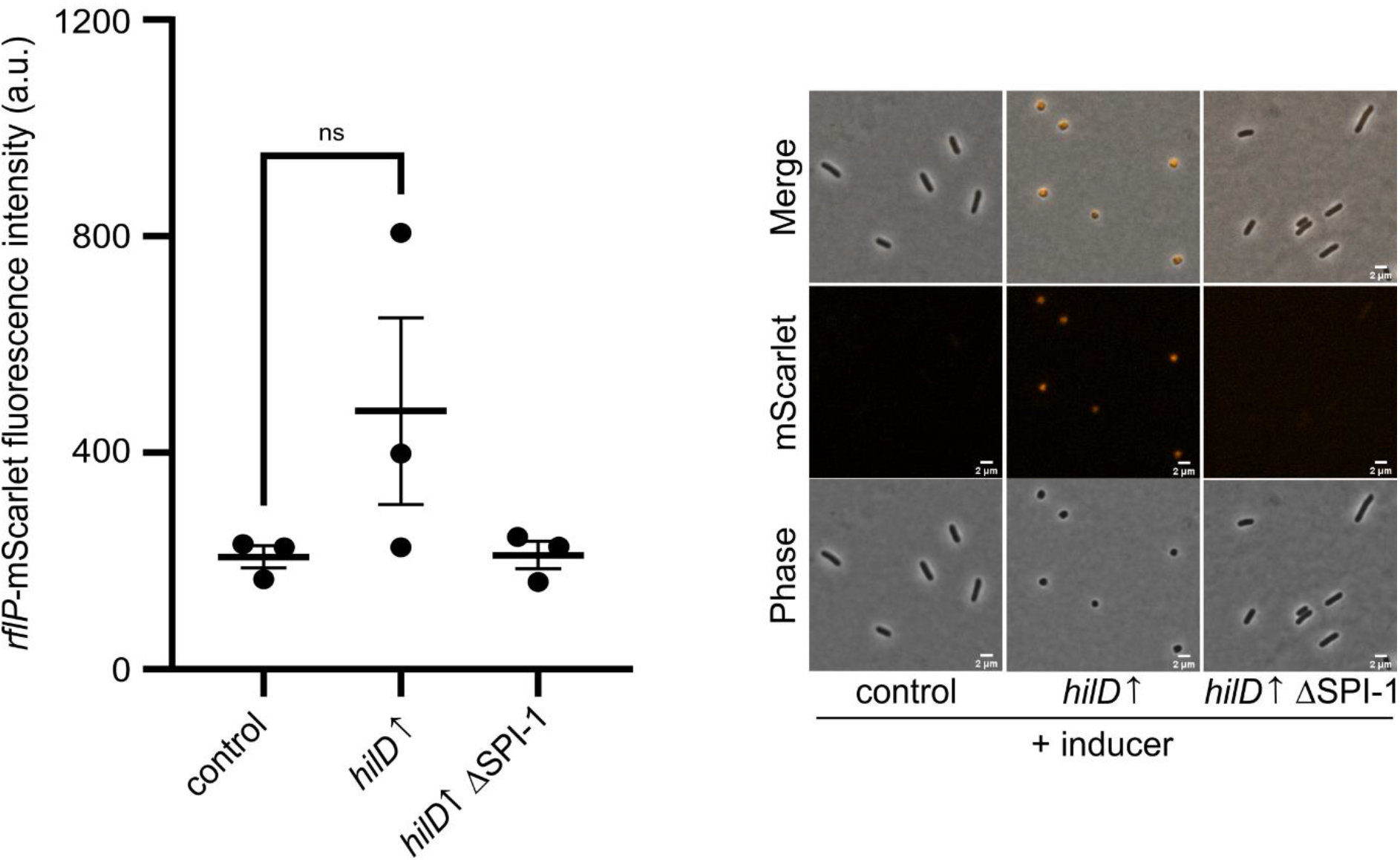
*rflP*-mScarlet expression under HilD inducing conditions. Strains were grown in presence of AnTc to induce HilD overproduction. Fluorescence intensities of *rflP*-mScarlet translational fusion were quantified in single-cells (left). Representative microscopy images are shown (right). Scale bar is 2 µm. Biological replicates are shown as individual data points. Horizontal bars represent the calculated mean of biological replicates. Error bars represent standard error of the mean. Statistical significances were determined using a two-tailed Student’s t-test (ns, *P* > 0.05). Strains analysed were EM13097 (control), EM13278 (*hilD*↑) and EM13363.(*hilD*↑ ΔSPI-1). *hilD*↑: strain expressing *hilD* under an inducible promoter. AnTc: anhydrotetracycline. a.u.: arbitrary units.

**Figure S6:**
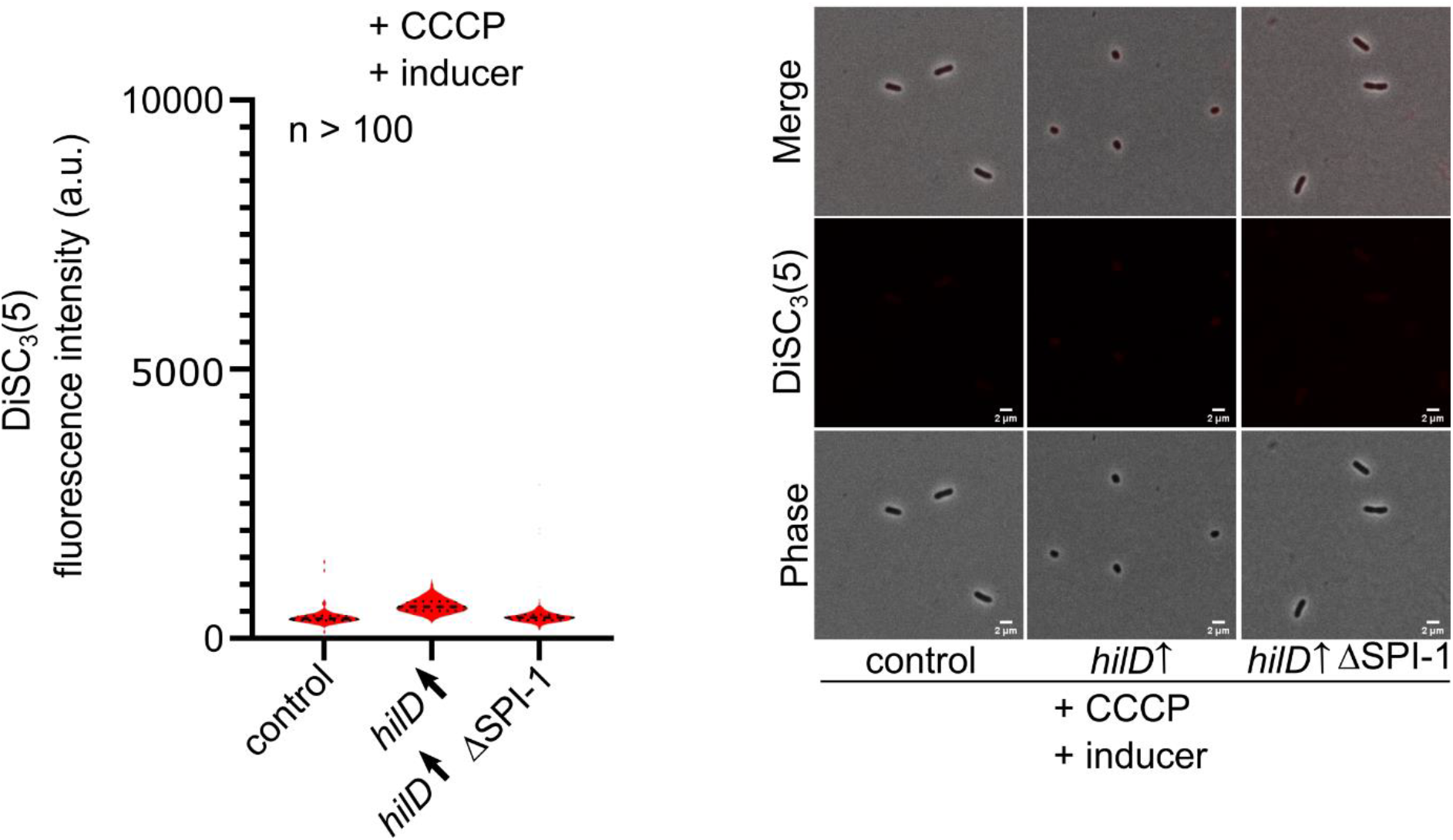
DiSC_3_(5) responds to CCCP-induced membrane depolarization. Strains were grown under HilD inducing conditions using AnTc followed by treatment with CCCP. Cells were then stained with the membrane potential sensitive dye DiSC_3_(5) and single-cell fluorescence intensities were quantified (left). Representative microscopy images are shown (right). Scale bar is 2 µm. Strains analysed were TH437 (control), TH17114 (*hilD*↑) and EM12479.(*hilD*↑ ΔSPI-1).*hilD*↑: strain expressing *hilD* under an inducible promoter. AnTc: anhydrotetracycline. a.u.: arbitrary units.

**Table S1. *Salmonella enterica* serovar Typhimurium strains used in this study**.

**Table S2. Plasmids used in this study**.

**Table S3. Primers used in this study**.

**Table S4. Transcriptional profile upon HilD induction in transcript per million (TPM)**

## References

1. Eng S-K, Pusparajah P, Ab Mutalib N-S, Ser H-L, Chan K-G, Lee L-H. Salmonella: A review on pathogenesis, epidemiology and antibiotic resistance. Front Life Sci. 2015;8: 284–293. doi:10.1080/21553769.2015.1051243

2. dos Santos AMP, Ferrari RG, Conte-Junior CA. Virulence Factors in Salmonella Typhimurium: The Sagacity of a Bacterium. Curr Microbiol. 2019;76: 762–773. doi:10.1007/s00284-018-1510-4

3. Hughes KT, Erhardt M. Bacterial Flagella. eLS. John Wiley & Sons, Ltd; 2011. doi:10.1002/9780470015902.a0000301.pub2

4. Wolfson EB, Elvidge J, Tahoun A, Gillespie T, Mantell J, McAteer SP, et al. The interaction of Escherichia coli O157 :H7 and Salmonella Typhimurium flagella with host cell membranes and cytoskeletal components. Microbiology. 166: 947–965. doi:10.1099/mic.0.000959

5. Horstmann JA, Lunelli M, Cazzola H, Heidemann J, Kühne C, Steffen P, et al. Methylation of Salmonella Typhimurium flagella promotes bacterial adhesion and host cell invasion. Nat Commun. 2020;11: 2013. doi:10.1038/s41467-020-15738-3

6. Yamaguchi T, Toma S, Terahara N, Miyata T, Ashihara M, Minamino T, et al. Structural and Functional Comparison of Salmonella Flagellar Filaments Composed of FljB and FliC. Biomolecules. 2020;10: 246. doi:10.3390/biom10020246

7. Erhardt M, Mertens ME, Fabiani FD, Hughes KT. ATPase-Independent Type-III Protein Secretion in Salmonella enterica. PLOS Genet. 2014;10: e1004800. doi:10.1371/journal.pgen.1004800

8. Mouslim C, Hughes KT. The Effect of Cell Growth Phase on the Regulatory Cross-Talk between Flagellar and Spi1 Virulence Gene Expression. PLOS Pathog. 2014;10: e1003987. doi:10.1371/journal.ppat.1003987

9. Yanagihara S, Iyoda S, Ohnishi K, Iino T, Kutsukake K. Structure and transcriptional control of the flagellar master operon of Salmonella typhimurium. Genes Genet Syst. 1999;74: 105–111. doi:10.1266/ggs.74.105

10. Chilcott GS, Hughes KT. Coupling of Flagellar Gene Expression to Flagellar Assembly in Salmonella enterica Serovar Typhimurium and Escherichia coli. Microbiol Mol Biol Rev. 2000;64: 694–708. doi:10.1128/MMBR.64.4.694-708.2000

11. Wang S, Fleming RT, Westbrook EM, Matsumura P, McKay DB. Structure of the Escherichia coli FlhDC Complex, a Prokaryotic Heteromeric Regulator of Transcription. J Mol Biol. 2006;355: 798–808. doi:10.1016/j.jmb.2005.11.020

12. Claret L, Hughes C. Functions of the subunits in the FlhD2C2 transcriptional master regulator of bacterial flagellum biogenesis and swarming11Edited by I. B. Holland. J Mol Biol. 2000;303: 467–478. doi:10.1006/jmbi.2000.4149

13. Hughes KT, Gillen KL, Semon MJ, Karlinsey JE. Sensing Structural Intermediates in Bacterial Flagellar Assembly by Export of a Negative Regulator. Science. 1993;262: 1277–1280. doi:10.1126/science.8235660

14. Chadsey MS, Karlinsey JE, Hughes KT. The flagellar anti-σ factor FlgM actively dissociates Salmonella typhimurium σ28 RNA polymerase holoenzyme. Genes Dev. 1998;12: 3123–3136. doi:10.1101/gad.12.19.3123

15. Wagner S, Grin I, Malmsheimer S, Singh N, Torres-Vargas CE, Westerhausen S. Bacterial type III secretion systems: a complex device for the delivery of bacterial effector proteins into eukaryotic host cells. FEMS Microbiol Lett. 2018;365. doi:10.1093/femsle/fny201

16. Lara-Tejero M, Galán JE. The Injectisome, a Complex Nanomachine for Protein Injection into Mammalian Cells. EcoSal Plus. 2019;8: 10.1128/ecosalplus.ESP-0039–2018. doi:10.1128/ecosalplus.ESP-0039-2018

17. Hayward RD, Koronakiss V. Direct modulation of the host cell cytoskeleton by Salmonella actin-binding proteins. Trends Cell Biol. 2002;12: 15–20. doi:10.1016/S0962-8924(01)02183-3

18. Lorkowski M, Felipe-López A, Danzer CA, Hansmeier N, Hensel M. Salmonella enterica Invasion of Polarized Epithelial Cells Is a Highly Cooperative Effort. Infect Immun. 2014;82: 2657–2667. doi:10.1128/IAI.00023-14

19. Cirillo DM, Valdivia RH, Monack DM, Falkow S. Macrophage-dependent induction of the Salmonella pathogenicity island 2 type III secretion system and its role in intracellular survival. Mol Microbiol. 1998;30: 175–188. doi:10.1046/j.1365-2958.1998.01048.x

20. Hensel M, Shea JE, Waterman SR, Mundy R, Nikolaus T, Banks G, et al. Genes encoding putative effector proteins of the type III secretion system of Salmonella pathogenicity island 2 are required for bacterial virulence and proliferation in macrophages. Mol Microbiol. 1998;30: 163–174. doi:10.1046/j.1365-2958.1998.01047.x

21. Ellermeier CD, Ellermeier JR, Slauch JM. HilD, HilC and RtsA constitute a feed forward loop that controls expression of the SPI1 type three secretion system regulator hilA in Salmonella enterica serovar Typhimurium. Mol Microbiol. 2005;57: 691–705. doi:10.1111/j.1365-2958.2005.04737.x

22. Saini S, Ellermeier JR, Slauch JM, Rao CV. The Role of Coupled Positive Feedback in the Expression of the SPI1 Type Three Secretion System in Salmonella. PLOS Pathog. 2010;6: e1001025. doi:10.1371/journal.ppat.1001025

23. Bajaj V, Hwang C, Lee CA. hilA is a novel ompR/toxR family member that activates the expression of Salmonella typhimurium invasion genes. Mol Microbiol. 1995;18: 715– 727. doi:10.1111/j.1365-2958.1995.mmi_18040715.x

24. Lostroh CP, Lee CA. The HilA Box and Sequences outside It Determine the Magnitude of HilA-Dependent Activation of PprgH from Salmonella Pathogenicity Island 1. J Bacteriol. 2001;183: 4876–4885. doi:10.1128/JB.183.16.4876-4885.2001

25. Keersmaecker S, Marchal K, Verhoeven T, Engelen K, Vanderleyden J, Detweiler C. Microarray Analysis and Motif Detection Reveal New Targets of the Salmonella enterica Serovar Typhimurium HilA Regulatory Protein, Including hilA Itself. J Bacteriol. 2005;187: 4381–91. doi:10.1128/JB.187.13.4381-4391.2005

26. Darwin KH, Miller VL. Type III secretion chaperone-dependent regulation: activation of virulence genes by SicA and InvF in Salmonella typhimurium. EMBO J. 2001;20: 1850– 1862. doi:10.1093/emboj/20.8.1850

27. Abby SS, Rocha EPC. The Non-Flagellar Type III Secretion System Evolved from the Bacterial Flagellum and Diversified into Host-Cell Adapted Systems. PLOS Genet. 2012;8: e1002983. doi:10.1371/journal.pgen.1002983

28. Desvaux M, Hébraud M, Henderson IR, Pallen MJ. Type III secretion: what’s in a name? Trends Microbiol. 2006;14: 157–160. doi:10.1016/j.tim.2006.02.009

29. Schavemaker PE, Lynch M. Flagellar energy costs across the tree of life. Michelot A, Walczak AM, Loiseau E, editors. eLife. 2022;11: e77266. doi:10.7554/eLife.77266

30. Paul K, Erhardt M, Hirano T, Blair DF, Hughes KT. Energy source of flagellar type III secretion. Nature. 2008;451: 489–492. doi:10.1038/nature06497

31. Wilharm G, Lehmann V, Krauss K, Lehnert B, Richter S, Ruckdeschel K, et al. Yersinia enterocolitica Type III Secretion Depends on the Proton Motive Force but Not on the Flagellar Motor Components MotA and MotB. Infect Immun. 2004;72: 4004–4009. doi:10.1128/IAI.72.7.4004-4009.2004

32. Minamino T, Namba K. Distinct roles of the FliI ATPase and proton motive force in bacterial flagellar protein export. Nature. 2008;451: 485–488. doi:10.1038/nature06449

33. Akeda Y, Galán JE. Chaperone release and unfolding of substrates in type III secretion. Nature. 2005;437: 911–915. doi:10.1038/nature03992

34. Kage H, Takaya A, Ohya M, Yamamoto T. Coordinated Regulation of Expression of Salmonella Pathogenicity Island 1 and Flagellar Type III Secretion Systems by ATP-Dependent ClpXP Protease. J Bacteriol. 2008;190: 2470–2478. doi:10.1128/JB.01385-07

35. Kühne C, Singer HM, Grabisch E, Codutti L, Carlomagno T, Scrima A, et al. RflM mediates target specificity of the RcsCDB phosphorelay system for transcriptional repression of flagellar synthesis in Salmonella enterica. Mol Microbiol. 2016;101: 841– 855. doi:10.1111/mmi.13427

36. Ellermeier CD, Slauch JM. RtsA and RtsB Coordinately Regulate Expression of the Invasion and Flagellar Genes in Salmonella enterica Serovar Typhimurium. J Bacteriol. 2003;185: 5096–5108. doi:10.1128/JB.185.17.5096-5108.2003

37. Spöring I, Felgner S, Preuße M, Eckweiler D, Rohde M, Häussler S, et al. Regulation of Flagellum Biosynthesis in Response to Cell Envelope Stress in Salmonella enterica Serovar Typhimurium. mBio. 9: e00736–17. doi:10.1128/mBio.00736-17

38. Wada T, Morizane T, Abo T, Tominaga A, Inoue-Tanaka K, Kutsukake K. EAL Domain Protein YdiV Acts as an Anti-FlhD4C2 Factor Responsible for Nutritional Control of the Flagellar Regulon in Salmonella enterica Serovar Typhimurium. J Bacteriol. 2011;193: 1600–1611. doi:10.1128/JB.01494-10

39. Petrone BL, Stringer AM, Wade JT. Identification of HilD-Regulated Genes in Salmonella enterica Serovar Typhimurium. J Bacteriol. 2014;196: 1094–1101. doi:10.1128/JB.01449-13

40. Banda MM, Zavala-Alvarado C, Pérez-Morales D, Bustamante VH. SlyA and HilD Counteract H-NS-Mediated Repression on the ssrAB Virulence Operon of Salmonella enterica Serovar Typhimurium and Thus Promote Its Activation by OmpR. J Bacteriol. 2019;201: e00530–18. doi:10.1128/JB.00530-18

41. Martínez LC, Banda MM, Fernández-Mora M, Santana FJ, Bustamante VH. HilD Induces Expression of Salmonella Pathogenicity Island 2 Genes by Displacing the Global Negative Regulator H-NS from ssrAB. J Bacteriol. 2014;196: 3746–3755. doi:10.1128/JB.01799-14

42. Olekhnovich IN, Kadner RJ. Role of Nucleoid-Associated Proteins Hha and H-NS in Expression of Salmonella enterica Activators HilD, HilC, and RtsA Required for Cell Invasion. J Bacteriol. 2007;189: 6882–6890. doi:10.1128/JB.00905-07

43. Bustamante VH, Martínez LC, Santana FJ, Knodler LA, Steele-Mortimer O, Puente JL. HilD-mediated transcriptional cross-talk between SPI-1 and SPI-2. Proc Natl Acad Sci. 2008;105: 14591–14596. doi:10.1073/pnas.0801205105

44. Gerlach RG, Jäckel D, Geymeier N, Hensel M. Salmonella Pathogenicity Island 4-Mediated Adhesion Is Coregulated with Invasion Genes in Salmonella enterica. Infect Immun. 2007;75: 4697–4709. doi:10.1128/IAI.00228-07

45. Main-Hester KL, Colpitts KM, Thomas GA, Fang FC, Libby SJ. Coordinate Regulation of Salmonella Pathogenicity Island 1 (SPI1) and SPI4 in Salmonella enterica Serovar Typhimurium. Infect Immun. 2008;76: 1024–1035. doi:10.1128/IAI.01224-07

46. Knodler LA, Celli J, Hardt W-D, Vallance BA, Yip C, Finlay BB. Salmonella effectors within a single pathogenicity island are differentially expressed and translocated by separate type III secretion systems. Mol Microbiol. 2002;43: 1089–1103. doi:10.1046/j.1365-2958.2002.02820.x

47. Navarre WW, Porwollik S, Wang Y, McClelland M, Rosen H, Libby SJ, et al. Selective Silencing of Foreign DNA with Low GC Content by the H-NS Protein in Salmonella. Science. 2006;313: 236–238.

48. Singer HM, Kühne C, Deditius JA, Hughes KT, Erhardt M. The Salmonella Spi1 Virulence Regulatory Protein HilD Directly Activates Transcription of the Flagellar Master Operon flhDC. J Bacteriol. 2014;196: 1448–1457. doi:10.1128/JB.01438-13

49. Schechter LM, Damrauer SM, Lee CA. Two AraC/XylS family members can independently counteract the effect of repressing sequences upstream of the hilA promoter. Mol Microbiol. 1999;32: 629–642. doi:10.1046/j.1365-2958.1999.01381.x

50. Boddicker JD, Knosp BM, Jones BD. Transcription of the Salmonella invasion gene activator, hilA, requires HilD activation in the absence of negative regulators. J Bacteriol. 2003;185: 525–533. doi:10.1128/JB.185.2.525-533.2003

51. Hurtado-Escobar GA, Grépinet O, Raymond P, Abed N, Velge P, Virlogeux-Payant I. H-NS is the major repressor of Salmonella Typhimurium Pef fimbriae expression. Virulence. 2019;10: 849–867. doi:10.1080/21505594.2019.1682752

52. Wang P, Robert L, Pelletier J, Dang WL, Taddei F, Wright A, et al. Robust growth of Escherichia coli. Curr Biol CB. 2010;20: 1099–1103. doi:10.1016/j.cub.2010.04.045

53. Sturm A, Heinemann M, Arnoldini M, Benecke A, Ackermann M, Benz M, et al. The Cost of Virulence: Retarded Growth of Salmonella Typhimurium Cells Expressing Type III Secretion System 1. PLOS Pathog. 2011;7: e1002143. doi:10.1371/journal.ppat.1002143

54. Schreiber G, Metzger S, Aizenman E, Roza S, Cashel M, Glaser G. Overexpression of the relA gene in Escherichia coli. J Biol Chem. 1991;266: 3760–3767. doi:10.1016/S0021-9258(19)67860-9

55. Gropp M, Strausz Y, Gross M, Glaser G. Regulation of Escherichia coli RelA Requires Oligomerization of the C-Terminal Domain. J Bacteriol. 2001;183: 570–579. doi:10.1128/JB.183.2.570-579.2001

56. Lee P-C, Zmina SE, Stopford CM, Toska J, Rietsch A. Control of type III secretion activity and substrate specificity by the cytoplasmic regulator PcrG. Proc Natl Acad Sci U S A. 2014;111: E2027–2036. doi:10.1073/pnas.1402658111

57. Gabel CV, Berg HC. The speed of the flagellar rotary motor of Escherichia coli varies linearly with protonmotive force. Proc Natl Acad Sci. 2003;100: 8748–8751. doi:10.1073/pnas.1533395100

58. Fung DC, Berg HC. Powering the flagellar motor of Escherichia coli with an external voltage source. Nature. 1995;375: 809–812. doi:10.1038/375809a0

59. Buttress JA, Halte M, Winkel JD te, Erhardt M, Popp PF, Strahl H. A guide for membrane potential measurements in Gram-negative bacteria using voltage-sensitive dyes. 2022; 2022.04.30.490130. doi:10.1101/2022.04.30.490130

60. Verstraeten N, Knapen WJ, Kint CI, Liebens V, Van den Bergh B, Dewachter L, et al. Obg and Membrane Depolarization Are Part of a Microbial Bet-Hedging Strategy that Leads to Antibiotic Tolerance. Mol Cell. 2015;59: 9–21. doi:10.1016/j.molcel.2015.05.011

61. Frye J, Karlinsey JE, Felise HR, Marzolf B, Dowidar N, McClelland M, et al. Identification of New Flagellar Genes of Salmonella enterica Serovar Typhimurium. J Bacteriol. 2006;188: 2233–2243. doi:10.1128/JB.188.6.2233-2243.2006

62. Colgan AM, Kröger C, Diard M, Hardt W-D, Puente JL, Sivasankaran SK, et al. The Impact of 18 Ancestral and Horizontally-Acquired Regulatory Proteins upon the Transcriptome and sRNA Landscape of Salmonella enterica serovar Typhimurium. PLOS Genet. 2016;12: e1006258. doi:10.1371/journal.pgen.1006258

63. Cooper KG, Chong A, Kari L, Jeffrey B, Starr T, Martens C, et al. Regulatory protein HilD stimulates Salmonella Typhimurium invasiveness by promoting smooth swimming via the methyl-accepting chemotaxis protein McpC. Nat Commun. 2021;12: 348. doi:10.1038/s41467-020-20558-6

64. Smith C, Stringer AM, Mao C, Palumbo MJ, Wade JT. Mapping the Regulatory Network for Salmonella enterica Serovar Typhimurium Invasion. mBio. 2016;7: e01024–16. doi:10.1128/mBio.01024-16

65. Morphological characterization of small cells resulting from nutrient starvation of a psychrophilic marine vibrio. [cited 6 Sep 2022]. doi:10.1128/aem.32.4.617-622.1976

66. Fida TT, Moreno-Forero SK, Heipieper HJ, Springael D 2013. Physiology and transcriptome of the polycyclic aromatic hydrocarbon-degrading Sphingomonas sp. LH128 after long-term starvation. Microbiology. 159: 1807–1817. doi:10.1099/mic.0.065870-0

67. James GA, Korber DR, Caldwell DE, Costerton JW. Digital image analysis of growth and starvation responses of a surface-colonizing Acinetobacter sp. J Bacteriol. 1995;177: 907–915. doi:10.1128/jb.177.4.907-915.1995

68. Miyazaki R, Minoia M, Pradervand N, Sulser S, Reinhard F, Meer JR van der. Cellular Variability of RpoS Expression Underlies Subpopulation Activation of an Integrative and Conjugative Element. PLOS Genet. 2012;8: e1002818. doi:10.1371/journal.pgen.1002818

69. Goormaghtigh F, Fraikin N, Putrinš M, Hallaert T, Hauryliuk V, Garcia-Pino A, et al. Reassessing the Role of Type II Toxin-Antitoxin Systems in Formation of Escherichia coli Type II Persister Cells. mBio. 2018 [cited 25 Jan 2022]. doi:10.1128/mBio.00640-18

70. Wada T, Tanabe Y, Kutsukake K. FliZ Acts as a Repressor of the ydiV Gene, Which Encodes an Anti-FlhD4C2 Factor of the Flagellar Regulon in Salmonella enterica Serovar Typhimurium. J Bacteriol. 2011;193: 5191–5198. doi:10.1128/JB.05441-11

71. Wei X, Bauer WD. Starvation-Induced Changes in Motility, Chemotaxis, and Flagellation of Rhizobium meliloti. Appl Environ Microbiol. 1998;64: 1708–1714. doi:10.1128/AEM.64.5.1708-1714.1998

72. Gray DA, Dugar G, Gamba P, Strahl H, Jonker MJ, Hamoen LW. Extreme slow growth as alternative strategy to survive deep starvation in bacteria. Nat Commun. 2019;10: 890. doi:10.1038/s41467-019-08719-8

73. The expression of virulence genes increases membrane permeability and sensitivity to envelope stress in Salmonella Typhimurium. PLoS Biol 20(4): e3001608. https://doi.org/10.1371/journal.pbio.3001608

74. Misselwitz B, Barrett N, Kreibich S, Vonaesch P, Andritschke D, Rout S, et al. Near surface swimming of Salmonella Typhimurium explains target-site selection and cooperative invasion. PLoS Pathog. 2012;8: e1002810. doi:10.1371/journal.ppat.1002810

75. Horstmann JA, Zschieschang E, Truschel T, de Diego J, Lunelli M, Rohde M, et al. Flagellin phase-dependent swimming on epithelial cell surfaces contributes to productive Salmonella gut colonisation. Cell Microbiol. 2017;19: e12739. doi:10.1111/cmi.12739

76. Ferreira JL, Gao FZ, Rossmann FM, Nans A, Brenzinger S, Hosseini R, et al. γ-proteobacteria eject their polar flagella under nutrient depletion, retaining flagellar motor relic structures. PLOS Biol. 2019;17: e3000165. doi:10.1371/journal.pbio.3000165

77. Sanderson KE, Roth JR. Linkage map of Salmonella typhimurium, edition VII. Microbiol Rev. 1988;52: 485–532. doi:10.1128/mr.52.4.485-532.1988

78. Datsenko KA, Wanner BL. One-step inactivation of chromosomal genes in Escherichia coli K-12 using PCR products. Proc Natl Acad Sci. 2000;97: 6640–6645. doi:10.1073/pnas.120163297

79. Hoffmann S, Schmidt C, Walter S, Bender JK, Gerlach RG. Scarless deletion of up to seven methyl-accepting chemotaxis genes with an optimized method highlights key function of CheM in Salmonella Typhimurium. Mantis NJ, editor. PLOS ONE. 2017;12: e0172630. doi:10.1371/journal.pone.0172630

80. Berg S, Kutra D, Kroeger T, Straehle CN, Kausler BX, Haubold C, et al. ilastik: interactive machine learning for (bio)image analysis. Nat Methods. 2019;16: 1226–1232. doi:10.1038/s41592-019-0582-9

81. Schindelin J, Arganda-Carreras I, Frise E, Kaynig V, Longair M, Pietzsch T, et al. Fiji: an open-source platform for biological-image analysis. Nat Methods. 2012;9: 676–682. doi:10.1038/nmeth.2019

82. Tinevez J-Y, Perry N, Schindelin J, Hoopes GM, Reynolds GD, Laplantine E, et al. TrackMate: An open and extensible platform for single-particle tracking. Methods. 2017;115: 80–90. doi:10.1016/j.ymeth.2016.09.016

83. Erhardt M. Fluorescent Microscopy Techniques to Study Hook Length Control and Flagella Formation. In: Minamino T, Namba K, editors. The Bacterial Flagellum: Methods and Protocols. New York, NY: Springer; 2017. pp. 37–46. doi:10.1007/978-1-4939-6927-2_3

84. Ducret A, Quardokus EM, Brun YV. MicrobeJ, a tool for high throughput bacterial cell detection and quantitative analysis. Nat Microbiol. 2016;1: 16077. doi:10.1038/nmicrobiol.2016.77

85. Green MR, Sambrook J, Sambrook J. Molecular cloning: a laboratory manual. 4th ed. Cold Spring Harbor, N.Y: Cold Spring Harbor Laboratory Press; 2012.

86. Deditius JA, Felgner S, Spöring I, Kühne C, Frahm M, Rohde M, et al. Characterization of Novel Factors Involved in Swimming and Swarming Motility in Salmonella enterica Serovar Typhimurium. PLOS ONE. 2015;10: e0135351. doi:10.1371/journal.pone.0135351

87. Pfaffl MW. A new mathematical model for relative quantification in real-time RT–PCR. Nucleic Acids Res. 2001;29: e45. doi:10.1093/nar/29.9.e45

88. Vandesompele J, De Preter K, Pattyn F, Poppe B, Van Roy N, De Paepe A, et al. Accurate normalization of real-time quantitative RT-PCR data by geometric averaging of multiple internal control genes. Genome Biol. 2002;3: research0034.1. doi:10.1186/gb-2002-3-7-research0034

