## Supplemental tables S1, S2, S3 for "SPI-1 virulence gene expression modulates motility of *Salmonella* Typhimurium in a proton motive force- and adhesins-dependent manner"

**Table S1.** ***Salmonella enterica*** **serovar Typhimurium strains used in this study.**

| Strain | Relevant characteristics | Reference |
|---|---|---|
| TH437 | *S. enterica* serovar Typhimurium wild-type strain LT2 | John Roth |
| TH16339 | Δ*araBAD*1065::$hilD^{+}$ | (1) |
| TH17114 | $P_{hilD}$::*tetRA* (tetracycline inducible *hilD*) | This study |
| EM93 | Δ*araBAD*1065::$hilD^{+}$ Δ*invH-sprB*::FRT | This study |
| EM228 | Δ*hutI-H*::$P_{sicA}$-eGFP | This study |
| EM808 | Δ*araBAD*1005::FRT | (2) |
| EM831 | Δ*araBAD*1182::$hilD_{\Delta HTH}$ | This study |
| EM840 | Δ*sseA-ssaU*::FRT (ΔSPI-2) Δ*araBAD*1065::$hilD^{+}$ | This study |
| EM899 | Δ*hutI-H*::$P_{sicA}$-eGFP Δ*araBAD*1065::$hilD^{+}$ | This study |
| EM900 | Δ*hutI-H*::$P_{sicA}$-eGFP Δ*araBAD*1005::FRT | This study |
| EM930 | Δ*araBAD*1183::$hilA^{+}$ | This study |
| EM3050 | $P_{flhDC}$22343 (-598 to -554 A=C, T=G) (randomized HilD binding site) Δ*araBAD*1005::FRT | This study |
| EM3051 | $P_{flhDC}$22343 (-598 to -554 A=C, T=G) (randomized HilD binding site) Δ*araBAD*1065::$hilD^{+}$ | This study |
| EM3052 | $P_{flhDC}$22343 (-598 to -554 A=C, T=G) (randomized HilD binding site) Δ*araBAD*1183::$hilA^{+}$ | This study |
| EM3059 | Δ$P_{flhDC}$(-598 to -554) (ΔHilD binding site) Δ*araBAD*1005::FRT | This study |

| | | |
|---|---|---|
| EM3060 | Δ$P_{flhDC}$(-598 to -554) (ΔHilD binding site) Δ*araBAD*1065::*hilD*$^{+}$ | This study |
| EM3061 | Δ$P_{flhDC}$(-598 to -554) (ΔHilD binding site) Δ*araBAD*1183::*hilA*$^{+}$ | This study |
| EM12302 | Δ*hutI-H*::P*sicA*-eGFP $P_{hilD}$::*tetRA* | This study |
| EM12144 | LT2 / pWSK29 (ApR) | This study |
| EM12145 | LT2 / p4830 (pWSK29-*tetR* $P_{tetA}$::*csgBACEFG*, ApR) | This study |
| EM12146 | LT2 / p4393 (pWSK29-tetR $P_{tetA}$::*safABCD*, ApR) | This study |
| EM12147 | LT2 / p4394 (pWSK29-tetR $P_{tetA}$::*stdABCD*, ApR) | This study |
| EM12148 | LT2 / p4396 (pWSK29-tetR $P_{tetA}$::*pefACDEF*, ApR) | This study |
| EM12354 | $P_{hilD}$::*tetRA* Δ*siiABCDEF* (ΔSPI-4) | This study |
| EM12648 | SR11 Δ12 Δ*csgA*::FKF $P_{hilD}$::*tetRA* | This study |
| EM12802 | LT2 / pBSB268 (pBAD18-GFP, ApR) | This study |
| EM12803 | $P_{hilD}$::*tetRA* / pBSB268 (pBAD18-GFP, ApR) | This study |
| EM13065 | *rpoS*-SAGASA-mCherry (C-ter translational fusion) $P_{hilD}$::*tetRA* | This study |
| EM13097 | *fliN*23482-mVenusNB-SAGASA (after aaM1) *rflP*23517::mScarlet (after aaM1) | This study |
| EM13100 | Δ*invH-sprB*::FRT attP22::[$P_{hilD}$::*tetRA*] Δ*sseA-ssaU*::FRT (deletes SPI-2) | This study |
| EM13146 | attP22::[$P_{hilD}$::*tetRA*] Δ*invH-sprB*::FCF (deletes SPI-1) | This study |
| EM13226 | *rpoS*-SAGASA-mCherry (C-ter translational fusion) / pTrc99a-FF4 (ApR) | This study |

| | | |
|---|---|---|
| EM13227 | *rpoS*-SAGASA-mCherry (C-ter translational fusion) / pEM13227 (pTrc99a-FF4-*relA*(aa1-455), ApR) | This study |
| EM13276 | attP22::[$P_{hilD}$::*tetRA*] Δ*invH-sprB*::FRT (deletes SPI-1) *rpoS*-SAGASA-mCherry (C-ter translational fusion) | This study |
| EM13278 | *fliN*23482-mVenusNB-SAGASA (after aaM1) *rflP*23518-mScarlet (before STOP) $P_{hilD}$::*tetRA* | This study |
| EM13363 | *fliN*23482-mVenusNB-SAGASA (after aaM1) *rflP*23518-mScarlet (before STOP) attP22::[$P_{hilD}$::*tetRA*] Δ*invH-sprB*::FKF (deletes SPI-1) | This study |

### Table S2. Plasmids used in this study.

| **Plasmids** | **Relevant characteristics** | **Reference** |
|---|---|---|
| pWSK29 | low copy number cloning vector (ApR) | Michael Hensel |
| p4830 | pWSK29-*tetR* $P_{tetA}$::*csgBACEFG* (ApR) | (3) |
| p4393 | pWSK29-tetR $P_{tetA}$::*safABCD* (ApR) | (3) |
| p4394 | p4394 (pWSK29-tetR $P_{tetA}$::*stdABCD* (ApR) | (3) |
| p4396 | p4396 (pWSK29-tetR $P_{tetA}$::*pefACDEF* (ApR) | (3) |
| pBSB268 | pBAD18-GFP (ApR) | (4) |
| pTrc99a-FF4 | IPTG-inducible expression vector (ApR) | (5) |
| pEM13227 | pTrc99a-FF4-*relA*(aa1-455) (ApR) | This study |

### Table S3. Primers used in this study.

| Number | Name | Sequence |
|---|---|---|
| 9 | Ara117fwd | CCACATTGAATATTTGCACAGCG |
| 59 | 5'-ParaB-hilA_50C-fw | actgtttctccatacctgtttttctggatggagtaagacgATGCCACATTTTAATCCTGT |
| 60 | 3'-ParaD-hilA_49C-rv | tcatcaacgcgccccccatgggacgcgtttttagaggcaTTACCGTAATTTAATCAAGCG |
| 128 | 5'-Dhut-PsicA-FPs-fw | aatgtctatacaacccatgacaggcaagagagcgacaggaGCCGCGTAAGGCAGTAGCGA |
| 129 | 3'-Dhut-PsicA-eGFP-rv | tgggcaccaccccggtgaacagttcttcgcctttgctcatTACTTACTCCTGTTATCTGT |
| 481 | 3'-hilD-dHTH_50C-fw | atcaacgcgccccccatgggacgcgtttttagaggcattATGACGAAGATATAATGTTGT |
| 5049 | 5’-Dspi-4-KanSceI_fw | acaaaaacattttattcacaatgtaatatcaggagacaacAGGGTTTTCCCAGTCACGAC |
| 5097 | DsiiA-F-KanSceI_new_rv | aagcagtaccacctgataacagcgacaagcgctgcttattTGCTTCCGGCTCGTATGTTG |
| 5098 | DsiiA-F_clean-del_new_fw | AATACGTATGGTTATAACGC |
| 5099 | DsiiA-F_clean-del_new rv | aagcagtaccacctgataacagcgacaagcgctgcttattGTTGTCTCCTGATATTACAT |
| 5340 | 5’-attP-Ptet-hilD_fw | aatgcgaaggtcgtaggttcgactcctattatcggcaccaGGGATTCCTGATGAAAATAG |
| 5341 | 3’-attP-Ptet-hilD_rv | tttttgagaaatgaggttgtacataagtgattgatttagaGTTAATGCGCAGTCTGAATT |
| 5591 | 3’-XbaI-relA (aa1-455)_rv | ctagtctagatcattattaCAACTGATAGGTGAATGGCA |
| 5718 | 5’_SAGASA-mCherry_fw | agcgcgggtgctagcgcgGTTTCCAAGGGCGAGGAGGA |
| 5719 | 3’_mCherry_rv | TTATTATTTGTACAGCTCAT |
| 5720 | 5’_rpoS-SAGASA-mCh_fw | gcagacgcaggggctgaatatcgaagcgctgttccgcgagAGCGCGGGTGCTAGCGCGGT |
| 5721 | 3’_rpoS- mCh_rv | gccagtcgacagactggcctttttttgacaagggtacttaTTATTATTTGTACAGCTCAT |
| 5748 | 5'-NdeI-relA_fw | cgccatatgGTCGCGGTAAGAAGTGCACA |
| 5751 | 5’_rpoS-tetRA-beforestop_fw | gcagacgcaggggctgaatatcgaagcgctgttccgcgagTTAAGACCCACTTTCACATT |
| 5752 | 3’_rpoS-tetRA-beforestop_rv | gccagtcgacagactggcctttttttgacaagggtacttaCTAAGCACTTGTCTCCTG |

### References


1. Singer HM, Kühne C, Deditius JA, Hughes KT, Erhardt M. The *Salmonella* Spi1 Virulence Regulatory Protein HilD Directly Activates Transcription of the Flagellar Master Operon flhDC. J Bacteriol. 2014 Apr 1;196(7):1448–57.

2. Paradis G, Chevance FFV, Liou W, Renault TT, Hughes KT, Rainville S, et al. Variability in bacterial flagella re-growth patterns after breakage. Sci Rep. 2017 Apr 28;7(1):1282.

3. Hansmeier N, Miskiewicz K, Elpers L, Liss V, Hensel M, Sterzenbach T. Functional expression of the entire adhesiome of *Salmonella enterica* serotype Typhimurium. Sci Rep. 2017 Dec;7(1):10326.

4. Dongre M, Singh B, Aung KM, Larsson P, Miftakhova R, Persson K, et al. Flagella-mediated secretion of a novel *Vibrio cholerae* cytotoxin affecting both vertebrate and invertebrate hosts. Commun Biol. 2018 Jun 7;1(1):1–12.

5. Ohnishi K, Fan F, Schoenhals GJ, Kihara M, Macnab RM. The FliO, FliP, FliQ, and FliR proteins of *Salmonella* typhimurium: putative components for flagellar assembly. J Bacteriol. 1997 Oct;179(19):6092–9.
